# A transcriptional clock of the human pluripotency transition

**DOI:** 10.1101/2025.03.13.643129

**Authors:** Louis Coussement, Matteo Ciarchi, Aljona Kolmogorova, Tim De Meyer, Steffen Rulands, Wolf Reik, Maria Rostovskaya

## Abstract

Developmental timing differs strikingly between mammals. All embryonic and some placental lineages emerge from the pluripotent epiblast in a temporally defined sequence, lasting about two weeks in the human embryo, but only two days in mice. Moreover, the order of lineage segregation and gene expression differ between the species. We used human pluripotent stem cells to recapitulate this window of epiblast development *in vitro.* Simultaneous profiling of gene expression and chromatin accessibility in single cells revealed a robust, autonomous switch between cell states during this process. We reconstructed the integrative gene regulatory network (GRN) of transcription factors (TFs) and signalling molecules. Notably, this revealed a transcriptional cascade including temporally close positive and distant inhibitory connections. We suggest that this cascade acts as a transcriptional clock and governs the directionality, timing and intrinsic decisions during epiblast development. From individual gene interactions, we derived a mechanistic mathematical model of the transcriptional clock of pluripotency that closely reproduced gene expression dynamics and identified key regulatory connections. Moreover, our model revealed a bistable switch governing the transition, thus translating the GRN inference into an interpretable mechanism. Strikingly, the GRN model derived for humans predicted the acceleration of developmental timing in mice when initialised with mouse-specific expression patterns, yet still leading to a human-like expression state. Therefore, TF levels explain developmental timing, whereas the architecture of the network defines its trajectory. Together, our work provides novel conceptual insights into the intrinsic mechanisms of developmental transitions in the human embryo.

**HIGHLIGHTS:**

- The human pluripotency transition is guided by an intrinsic developmental programme
- A cascade of transcription factors defines directionality and timing of the pluripotency transition through positive and negative interactions
- Transcription factor levels explain developmental timing
- The architecture of the gene regulatory network defines its trajectory

**GRAPHICAL ABSTRACT:** 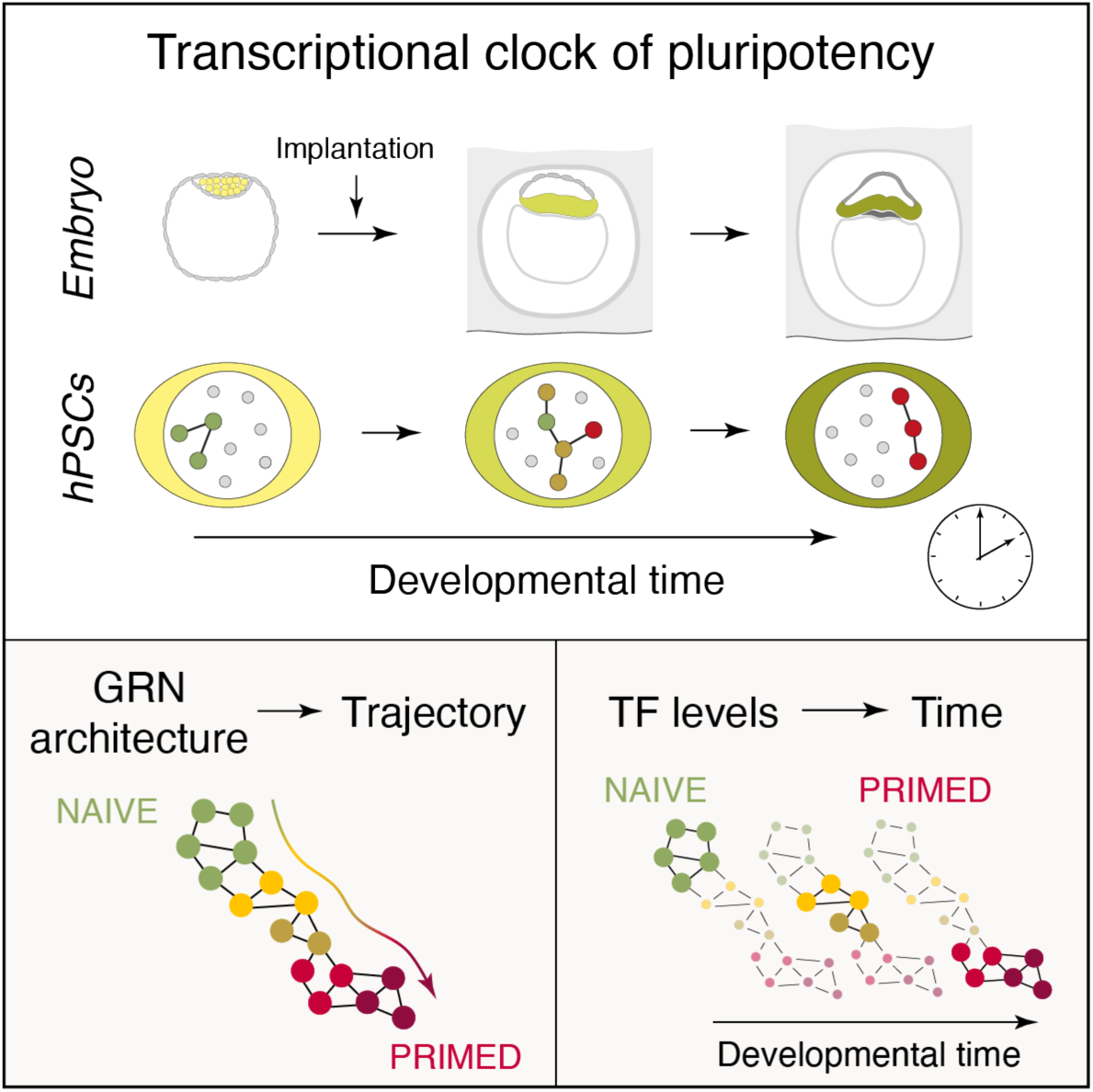

## INTRODUCTION

During development, cell differentiation follows a specific, reproducible timetable; however, the mechanisms tracking time remain largely unknown^1^. In mammals, the entire organism emerges from a small group of 10-20 equivalent cells called the epiblast. Differentiated lineages segregate from the pluripotent epiblast in a stereotypical, temporally defined sequence, over a period lasting for about 2 weeks in human. Surprisingly, the timing and order of lineage segregation vary considerably across species^2–4^, suggesting that this timetable maybe based on a simple molecular mechanism adjustable through evolution.

Timekeeping mechanisms have been identified in several developmental systems, including the mid-blastula transition in *Xenopus*^5^ and *Drosophila*^6,7^, larval progression in *C. elegans*^8^, somite formation^9^ and *Hox* genes activation^10^ in mammals. Changes in developmental timing, or heterochrony, often account for alterations in organ size, shape and complexity, as well as the overall body plan, serving as a major driving force in evolution. For example, in human compared to non-human primates, the window for neural progenitor expansion is extended, while the timing of cortical neurogenesis is delayed, which have been linked to the expansion and gyrification of the human cerebral cortex^11^. Thus, elucidating timekeeping mechanisms is essential for understanding critical developmental processes and differentiation pathways.

Several components are integral to the concept of developmental timing, including the order of events, the relative duration of stages, and their pace. Various molecular mechanisms have been proposed to regulate developmental timing, including protein stability^12^, rate of biochemical reactions^13^, mitochondrial metabolic activity^14,15^, and epigenetic pathways^16^. These mechanisms have been implicated in controlling the pace of motor neuron differentiation, neuronal maturation, and segmentation clock oscillations. They are not mutually exclusive and likely cooperate in regulating developmental timing^1^. At the same time, they govern cellular processes at a global level, likely changing their pace, but not tracking time increments or scheduling events. Nevertheless, the order of events is not always conserved across species (known as sequence heterochrony), suggesting that such global scaling alone is insufficient to explain temporal control. Therefore, these processes can be considered “pace-makers” that proportionally scale developmental timing across species, while time tracking is likely carried out by mechanistically distinct ”time-keepers”, potentially under “pace-maker” control.

Pluripotent stem cells represent *in vitro* counterparts of the embryonic epiblast^17–19^. Two distinct states of pluripotency, naïve and primed, correspond to the emergent pre-implantation and late pre-gastrulating epiblast, respectively^20^. In previous work, we developed a human pluripotent stem cell (hPSC)-based experimental system that recapitulates the entire 2-week long window of epiblast development, known as the pluripotent state transition, or the naïve-to-primed transition^21^.

The pluripotent state transition *in vitro* closely follows the molecular dynamics and timing of epiblast development^21,22^. Interestingly, developmental timing is generally preserved even outside the embryo context, as seen in motor neuron differentiation from mouse and human PSC *in vitro*^12,16^, oscillations of the segmentation clock in PSC-derived presomitic mesoderm *in vitro* ^23–25^, and the maturation of primate cortical neurons transplanted into the mouse brain^26–28^. Together, these findings strongly suggest that the timing of developmental programmes is governed by intrinsic mechanisms.

In this work, we reconstructed the molecular network of transcription factors (TFs) governing the pluripotent state transition using single-cell multiomics data. Through mathematical modelling and functional experiments, we demonstrated that this regulatory system acts as an intrinsic timer, based on a combination of positive and negative feedback loops and threshold behaviour. We found that the properties of this regulatory network explain the species-specific timetable of early embryogenesis. To our knowledge, this is the second developmental timer in human explained at the molecular level, since the discovery of the segmentation clock in 1997^9^.

## RESULTS

### Differences in developmental timing are associated with restructuring of the epiblast progression programme

To elucidate the mechanisms controlling developmental timing, we compared the molecular dynamics of epiblast development across species with markedly different paces. The pluripotent state transition, from the emergence of the epiblast in the pre-implantation blastocyst until the appearance of somatic lineages in the gastrulating embryo, takes 2-3 days in mice and 10-14 days in humans^2–4^. Moreover, the order of lineage segregation during this period does not match between these species (Figure S1A).

We compared the epiblast gene expression dynamics in mouse embryos *in utero* and human embryos cultured *in vitro* using published single-cell RNAseq datasets^29,30^ (Figure S1B, Table S1). Hierarchical clustering identified several stages during the epiblast progression in both mouse (pre-implantation, mPreEPI, and post-implantation, mPostEPI-E, -L1, -L2) and human (pre-implantation, hsPreEPI, and post-implantation, hsPostEPI-E1, -E2) embryos. Mouse and human epiblast expression patterns generally correlated (Figure S1C), indicating the overall similarity of the epiblast progression in both species.

As expected, the epiblast lineage was marked by *POU5F1*/*Pou5f1* in the absence of *GATA6*/*Gata6*^18,31^. In humans, the pre-implantation epiblast expressed *FBP1*, *KLF5*, and *TFCP2L1*, whilst the post-implantation cells were positive for *FZD7* and *SFRP2*^32,33^. In mice, the pre- and post-implantation stages were marked by *Fgf4*, *Klf2*, *Tfcp2l1,* and *Pou3f1, Zic5*, respectively^32^ (Figure S1D).

For each species, the dynamics of the most differentially expressed genes during the pluripotent state transition were similar between the epiblasts *in vivo* and pluripotent stem cells *in vitro* (Figure S1E and S1F). However, the expression changes of the most variable human epiblast genes did not align with their dynamics in mice. This discrepancy was also reflected in differences between hPSCs and mESCs (Figure S1G-S1I, Table S1). Only approximately 32.0% of downregulated and 61.5% of upregulated genes in humans showed the same direction of change in mice. This inconsistency was also observed among the most dynamic transcription factors (TFs) (Figure S1J).

Altogether, our analysis suggests that the key events, including gene expression and lineage segregation, occur in a different order during the pluripotency window in mouse and human. Therefore, the molecular programme governing epiblast development is not simply proportionally scaled among species with different developmental paces but is rather restructured.

### The pluripotency transition is guided by an autonomous molecular programme

The pluripotent state transition of hPSCs recapitulates the molecular dynamics and timing of human epiblast development^21,22,34^. To understand the molecular programme governing epiblast development, we performed simultaneous profiling of the transcriptome and chromatin accessibility in single cells in hPSCs during this transition using 10X technology (Figure 1A).

**Figure 1.**
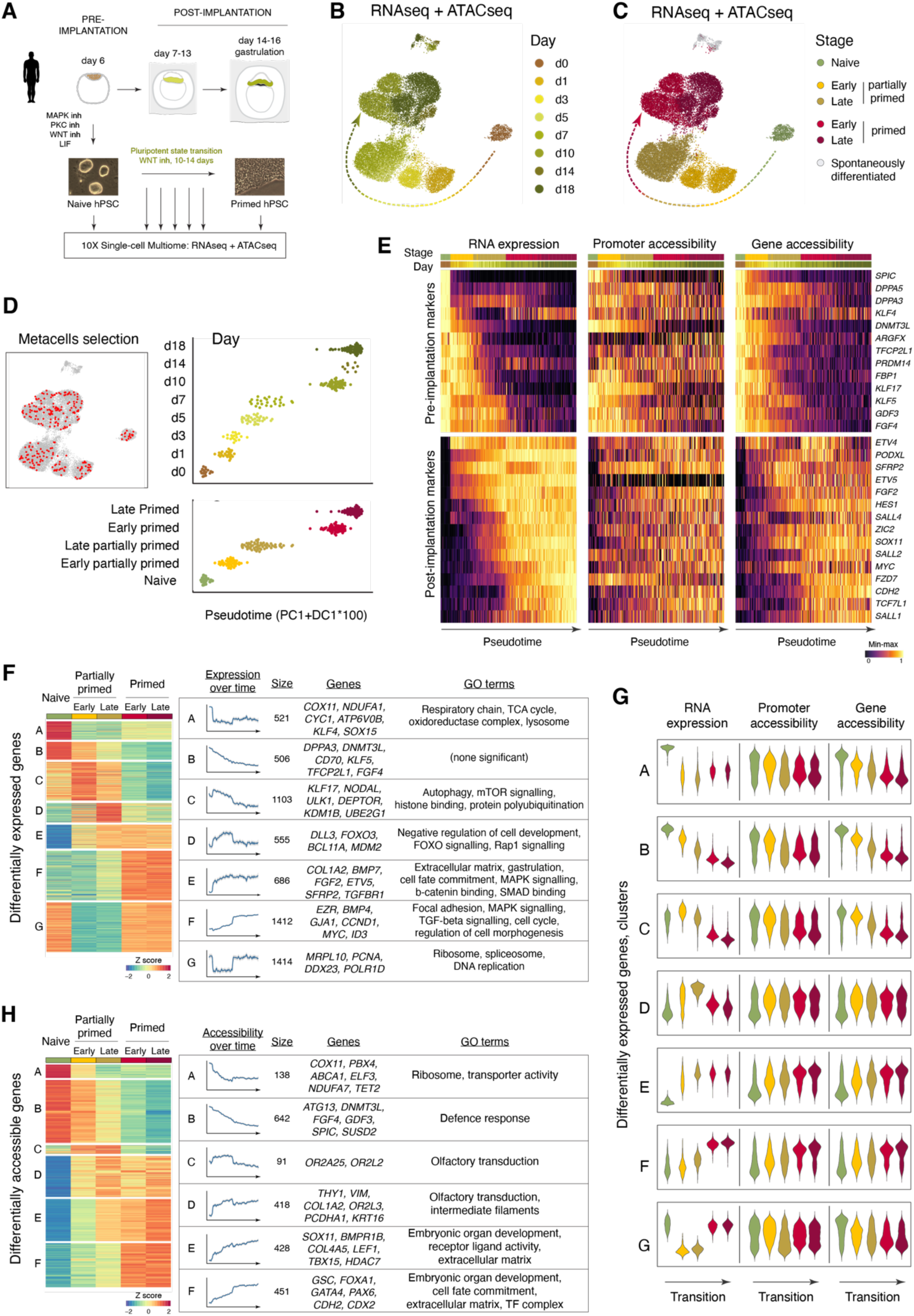
Simultaneous profiling of transcriptome and chromatin accessibility in single cells reveals the autonomous switch during the pluripotent state transition (also see Figures S2 and S3, Table S2). (A) Experimental setup. (B) UMAP based on MOFA, with colors indicating day of the transition. (C) UMAP based on MOFA, with colors indicating cell cluster. (D) Metacell selection. (E) Heatmaps showing RNA expression and chromatin accessibility of known pluripotency markers in metacells, ordered along pseudotime. (F) Global gene expression dynamics during the pluripotent state transition. (G)Correspondence between gene expression and chromatin accessibility in hPSCs. (H) Global chromatin accessibility dynamics in genic regions during the pluripotent state transition.

A total of 36,637 cells passed quality control (QC) for RNAseq (ranging from 907 to 8,192 cells per time point), with a median of 16,163 reads and 5,123 genes detected per cell (Figure S2A-S2I). A total of 27,701 cells passed QC for ATACseq (ranging from 904 to 6,067 cells per time point), with a median of 17,570 ATACseq fragments per cell (Figure S3A-S3F). A total of 23,738 cells passing QC for both RNAseq and ATACseq were used for the integration of data modalities using MOFA+^35^.

Dimensionality reduction analysis showed that the populations were relatively homogeneous at each time point, with only a small proportion of spontaneously differentiated cells (Figure 1B, S2F and S2G, S3G and S3H). The transition progressed directionally in a synchronous and coordinated manner. Interestingly, this analysis suggested that the transition was a stepwise rather than a gradual process, with two major switches and three cell states: naïve, partially primed and primed.

The first switch, from day 0 to day 1, could be explained by the change in culture conditions that initiated the transition. However, the second switch, from day 7 to day 10, occurred under constant conditions, indicating that it was not induced exogenously but rather driven by an autonomous decision to switch cell states. Once the primed state was established on day 10, it remained stable on day 14 and 18, thus the stepwise change between day 7 and 10 was unlikely to be due to the time gap in sampling.

For detailed analysis of the transition, we identified five temporal cell clusters – naïve, early and late partially primed, early and late primed – through hierarchical clustering based on RNAseq and ATACseq, either individually or in combination (Figure 1C, S2J and S3I). Next, we aggregated the data into metacells using SEACell^36^. After QC (for details, see STAR Methods), 250 metacells were selected, each representing an average of 88.9 cells. This approach overcomes the sparsity of single-cell data while retaining information about cell heterogeneity. Pseudotime analysis showed that the metacells aligned along the trajectory in the correct temporal order (Figure 1D).

During transition, pre-implantation epiblast markers were downregulated (*KLF4*, *TFCP2L1*, *ARGFX*, *DNMT3L*, *KLF5*, *KLF17*, *FGF4*, *SPIC*, *DPPA3*, *DPPA5*, *PRDM14*, *FBP1*, *GDF3*), while post-implantation markers were upregulated (*PODXL*, *ETV4*, *ETV5*, *SFRP2*, *FGF2*, *HES1*, *SOX11*, *SALL1*, *SALL2*, *SALL4*, *ZIC2*, *MYC*, *FZD7*, *CDH2*, *TCF7L1*) ^18,21,37^ (Figure 1E). We tested the accessibility of promoter regions (from 500bp upstream to 100bp downstream of the transcription start site, TSS) and gene loci (from 5,000bp upstream of the TSS to the gene terminator) of these epiblast markers. The accessibility of these gene loci followed the gene expression better than promoter accessibility alone.

The RNAseq analysis revealed 6,197 genes differentially expressed (DEGs) between any two time points (FDR <0.01, abs(log_2_(Fold Change)) > 1), out of 36,601 genes, forming 7 dynamic gene clusters (Figure 1F, Table S2). Supporting our global analysis of cell states, most changes occurred during the switch from naïve to partially primed hPSCs and from partially primed to primed hPSCs. The early wave of changes was associated with downregulation of genes involved in oxidative phosphorylation and TCA cycle, and with upregulation of extracellular matrix production, signalling factors, and developmental genes. The late wave of changes included downregulation of genes involved in autophagy and epigenetic regulation, while adhesion and signalling molecules, and regulators of morphogenesis were upregulated. We examined the accessibility of promoters and gene loci across the dynamically expressed gene clusters (Figure 1G). Consistent with previous reports^38^ and our analysis of pluripotency markers (Figure 1E), gene expression aligned more closely with gene accessibility than with promoter accessibility alone.

The ATACseq analysis showed 70,143 differentially accessible peaks (DAPs) from a total of 136,109 peaks and 2,168 differentially accessible genes (DAGs) from 24,919 genes (FDR <0.01, abs(log_2_(Fold Change)) > 1) (Figure S3J and 1H, Table S2).

There were more DAPs showing a loss of accessibility rather than a gain (40,030 vs 30,113, respectively); and a larger fraction of those DAPs located distally in the genome (43.2-46.0% vs 37.7-39.1%, respectively) (Figure S3K). This is consistent with the global chromatin compaction during the transition^22^. Hierarchical clustering revealed 6 DAG clusters, including developmental genes, extracellular matrix and signalling factors, and genes not expressed in hPSCs, such as olfactory receptors. Therefore, the dynamic chromatin state is associated with both, the gene expression changes and the global shifts in genome accessibility, during the transition.

Taken together, single-cell multiomics revealed that the pluripotency transition is governed by an autonomous, temporally defined molecular programme that incorporates a cell state switch.

### Signalling pathway dynamics during the pluripotent state transition

Our differential gene expression analysis suggested a shift in signalling pathway activities during the pluripotent state transition. To investigate these dynamics, we developed an approach combining the existing computational tools (Figure 2A).

**Figure 2.**
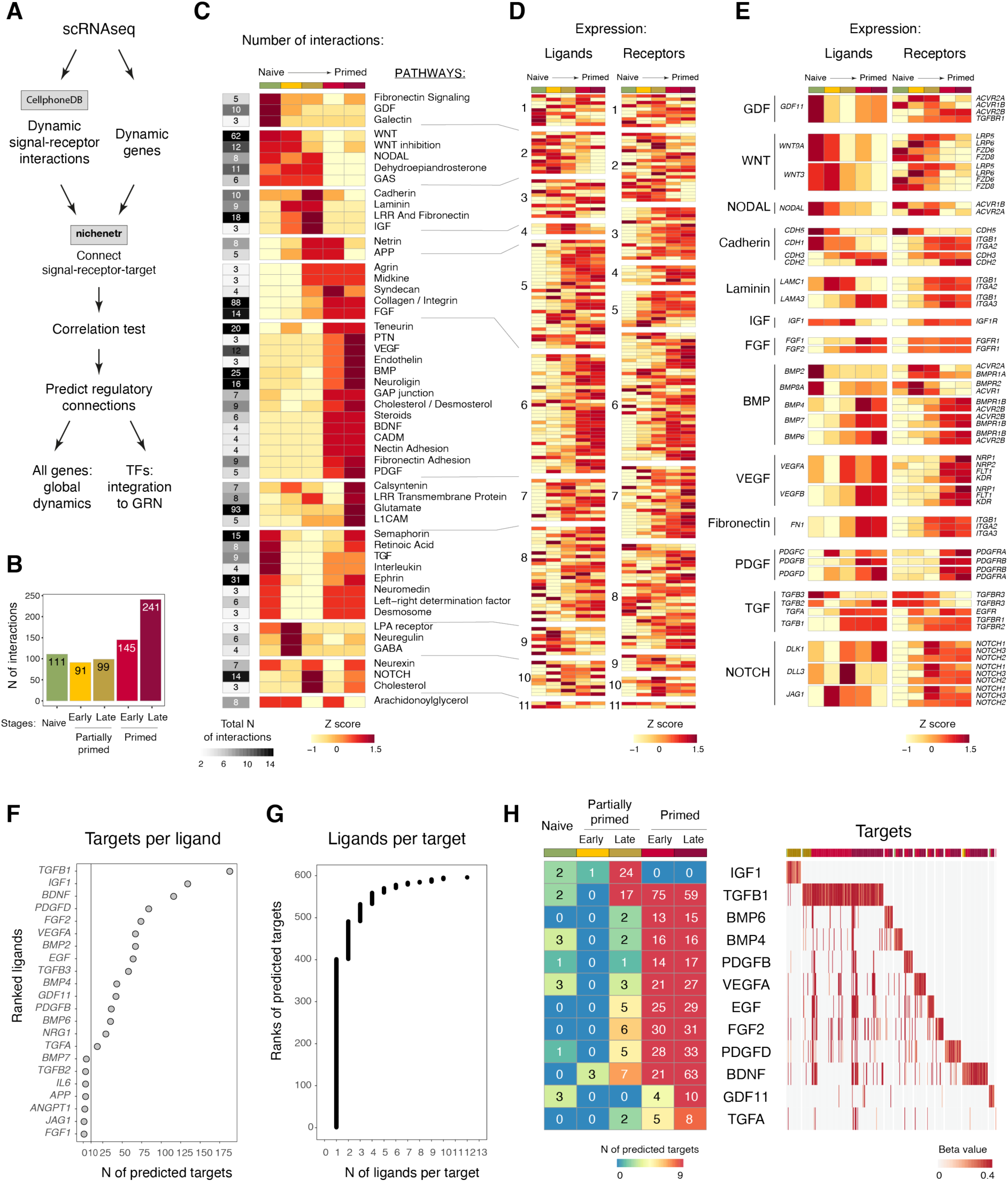
Signalling pathway dynamics during the pluripotent state transition (also see Table S3). (A) Analysis workflow: scRNAseq data were analysed to predict dynamic ligand-receptor pairs by CellPhoneDB, and to identify dynamic genes by differential expression analysis. These results were then integrated into the NicheNetR tool to predict ‘ligand-receptor-target’ combinations. Potential regulatory connections were identified based on expression correlations among predicted ‘ligand-receptor-target’ sets. (B) Bar plot showing the numbers of dynamic interactions per cell cluster, as predicted using CellPhoneDB. (C) Heatmap showing the numbers of dynamic interactions per pathway across cell clusters based on CellPhoneDB predictions. (D) Heatmap showing gene expression levels for ligands and receptors involved in dynamic interactions, as predicted using CellPhoneDB. (E) Examples of signalling molecules and their receptors involved in dynamic interactions, identified through CellPhoneDB predictions. (F) Dot plot showing the number of dynamic genes identified as targets of dynamic ligands, based on NicheNetR predictions. (G) Dot plot showing the number of ligands per target gene, as predicted by NicheNetR. (H) Predicted regulatory connections from signalling molecules to differentially expressed genes (left: numbers of DEGs per ligand across stages; right: linear regression coefficients between ligands and their putative targets). The putative target genes were associated to cell clusters according to their maximum expression.

Using CellPhoneDB^39–41^, we predicted dynamic ‘ligand-receptor’ pairs and observed an overall increase in the number of signalling interactions during the transition (Figure 2B, Table S3). Notably, while WNT, GDF, NODAL showed a decrease in the number of interactions, FGF, BMP, collagen, VEGF pathways exhibited an increase. More complex dynamics was observed within the TGF pathway, where specific ligand-receptor pairs, such as TGFB3-TGFBR3, were exchanged for other combinations like TGFB1-TGFBR1 (Figure 2C-2E).

Next, we employed NicheNetR^42^ to connect the dynamic signals identified by the CellPhoneDB to their potential target genes. This tool infers ‘ligand-receptor-target’ connections and estimates the strength of these predicted regulatory links (or weights) based on prior knowledge. To further evaluate the regulatory potential of these ‘ligand-receptor-target’ combinations, we applied the following criteria: (1) evidence for the connection estimated through the weight; (2) the expression dynamics of a ligand and its putative target should correlate; (3) the receptor’s dynamics should not contradict that of the respective signal (i.e., the receptor should either change in the same direction or remain unchanged).

Using this approach, we predicted 22 ligands potentially regulating 596 DEGs during the transition (Figure 2F, Table S3), with 491 genes of these connected to one or two ligands (Figure 2G). TGFB1 and IGF1 were predicted to have the highest number of potential targets (188 and 134 targets, respectively), which is consistent with their established roles in regulating pluripotency^43–46^. Notably, the majority of the positively regulated putative targets of the dynamic signalling pathways were expressed at the primed stage (81.2%), and the greatest change in signalling activities coincided with the switch from the partially primed to the primed state (Figure 2H). Therefore, the pluripotent state transition is associated with a shift and overall increase of signalling pathway activities.

### Reconstruction of the integrative dynamic gene regulatory network operating during the pluripotent state transition

We aimed to reconstruct the autonomous molecular programme that drives the pluripotent state transition with defined timing and governs the cell state switches. We hypothesized that this programme represents an endogenous network of TFs connected through feedforward and negative feedback loops, potentially coordinated by auto- and paracrine signalling. To achieve this, we developed an approach to reconstruct this integrative gene regulatory network (GRN) of TFs and signals (Figure 3A).

**Figure 3.**
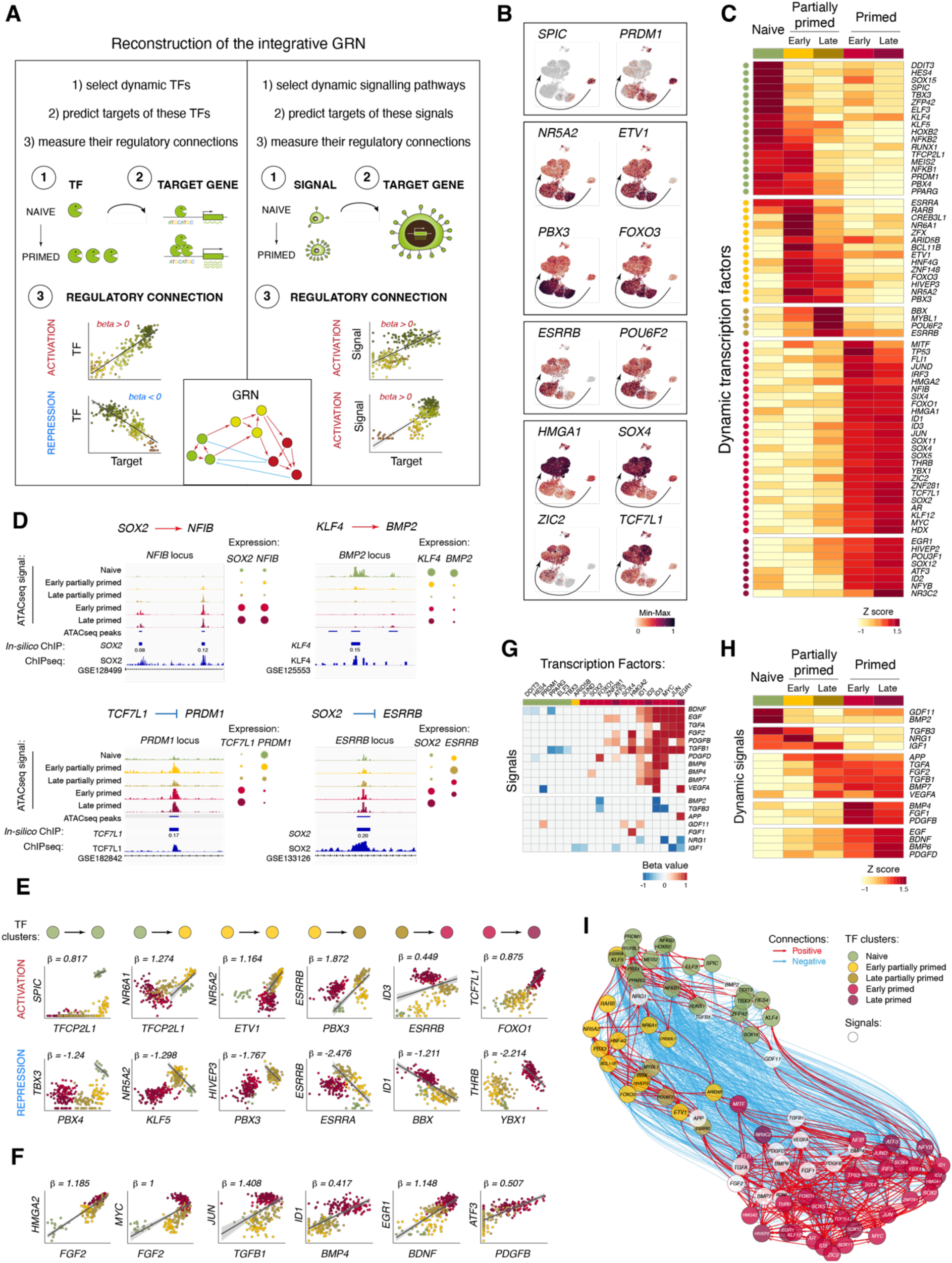
Reconstruction of the integrative dynamic gene regulatory network operating during the pluripotent state transition (also see Figures S4 and S5, Tables S4 and S5). (A) Schematic of the method. (B) Examples of TFs dynamically expressed during the pluripotent state transition. UMAPs based on scMultiome, with colors indicating relative expression. (C) Heatmap showing the most dynamically expressed TFs. (D) Genome browser tracks showing dynamic ATACseq peaks with predicted transcription factors binding sites. Experimental ChIPseq data are shown at the bottom, along with the predicted *in silico* ChIP binding sites and the corresponding scores. (E) Scatter plots showing examples of positive and negative associations between TFs expressed at different stages of the transition. Each dot represents a metacell colored by the corresponding stage of the transition. The line indicates a linear regression of TF expression, considering metacells used for calculations. (F) Scatter plots showing examples of associations between signalling molecules and TFs. (G) Heatmap showing significant correlations between dynamic signalling molecules and TFs. (H) Heatmap showing the expression levels of signalling molecules with predicted regulatory connections to the dynamic TFs. (I) UMAP representation of the GRN operating during the pluripotent state transition.

TFs were selected based on their upregulation in a given cell cluster (naïve, early and late partially primed, early and late primed) compared to at least 3 of the other 4 clusters. Thus, these TFs were either specific to one or shared between two stages of the transition. Non-unique TFs were associated with the earlier stage, resulting in a catalogue of 70 TFs that marked the 5 stages of the transition (Figure 3B and 3C, S4A and S4B). It included well-known naïve and primed pluripotency markers, such as *KLF4*, *KLF5*, *SOX11*, *ZIC2*, as well as less-characterised in the context of pluripotency factors like *RUNX1*, *PPARG*. The expression of these TFs was validated using an independent bulk RNAseq dataset^21^ and qRT-PCR (Figure S4C and S4D).

The network of TFs was reconstructed using a previously reported method^38^. First, we defined the putative cis-regulatory elements as ATACseq peaks within 50kb of the target gene body (Figure S5A and S5B). Most of the peaks were located distally relative to the genes (Figure S5C and S5D) and a large fraction displayed a positive correlation with the expression of their respective putative target genes (Figure S5E-S5G). Next, we predicted TF binding to these regulatory elements using an *in silico* ChIP approach. Each ‘TF-regulatory element’ combination was quantitatively assessed using an *in silico* ChIP score, which incorporated (1) the TF binding motif score, (2) the correlation between the chromatin accessibility of this region and the expression level of the TF, and (3) the average chromatin accessibility (Figure S5H). The approach was validated through comparisons to existing ChIPseq datasets (Figure S5I-S5K) enabling us to define an optimal *in silico* ChIP score threshold of 0.06. This resulted in a comprehensive *in silico* ChIP library comprising 781 TFs, their 2,086,946 potential target cis-regulatory elements (with a median of 1,680 regions per TF) and 32,255 unique target genes (with a median of 4,912 genes per TF) (Table S4). Finally, the TF-target gene pairs were quantitatively assessed for the strength of their regulatory connection through linear regression of their expression levels (beta-values) (Figure 3D and 3E, also see STAR Methods for details).

Next, we incorporated signalling molecules into the regulatory network. Our previous analysis (Figure 2H) identified 18 signals connected to 20 TFs of the network, with 13 of these TFs being expressed in the primed state (Figure 3F-3H). Regression coefficients provided quantitative measures of their potential connections. Notably most of these connections (62 out of 84) were activating.

Finally, we constructed a GRN of TFs and signals using linear regression coefficients (beta values) as measures of their regulatory connections (Figure 3I, Table S5). After removing 4 TFs (*ZFX*, *POU3F1*, *HDX*, *ZNF148*) and 2 signalling factors (*IGF1, EGF*) that lacked activating incoming connections, the network consisted of 66 TFs and 16 signals. All 82 network members sent the regulatory connections, with the total number of outgoing connections per member ranging from 1 to 50 (median 16) (Figure S5L and S5M). The activating connections were predominantly temporally short-range, targeting genes within the same or the next consecutive stage. In contrast, the repressing connections were broader in scope, targeting all clusters and including long-range connections across the entire transition. Notably, repressing connections were more abundant than activating ones (1,088 vs 389, respectively), likely due to the non-overlapping expression patterns of the GRN members, which made activating connections less frequent within this set.

Together, we reconstructed an integrative GRN of TFs and signals and hypothesised that this autonomous molecular mechanism directionally guides the pluripotency transition with defined timing.

### Transcription factor network acts as a timer of the transition

To understand how the GRN guides the pluripotent state transition, we created a mathematical model to quantitatively describe this process based on our multiomics data (Figure 4A). The model’s inputs included the initial gene expression levels of the GRN members and the strength of their regulatory connections to target genes, quantified through regression coefficients between their expression levels over time.

**Figure 4.**
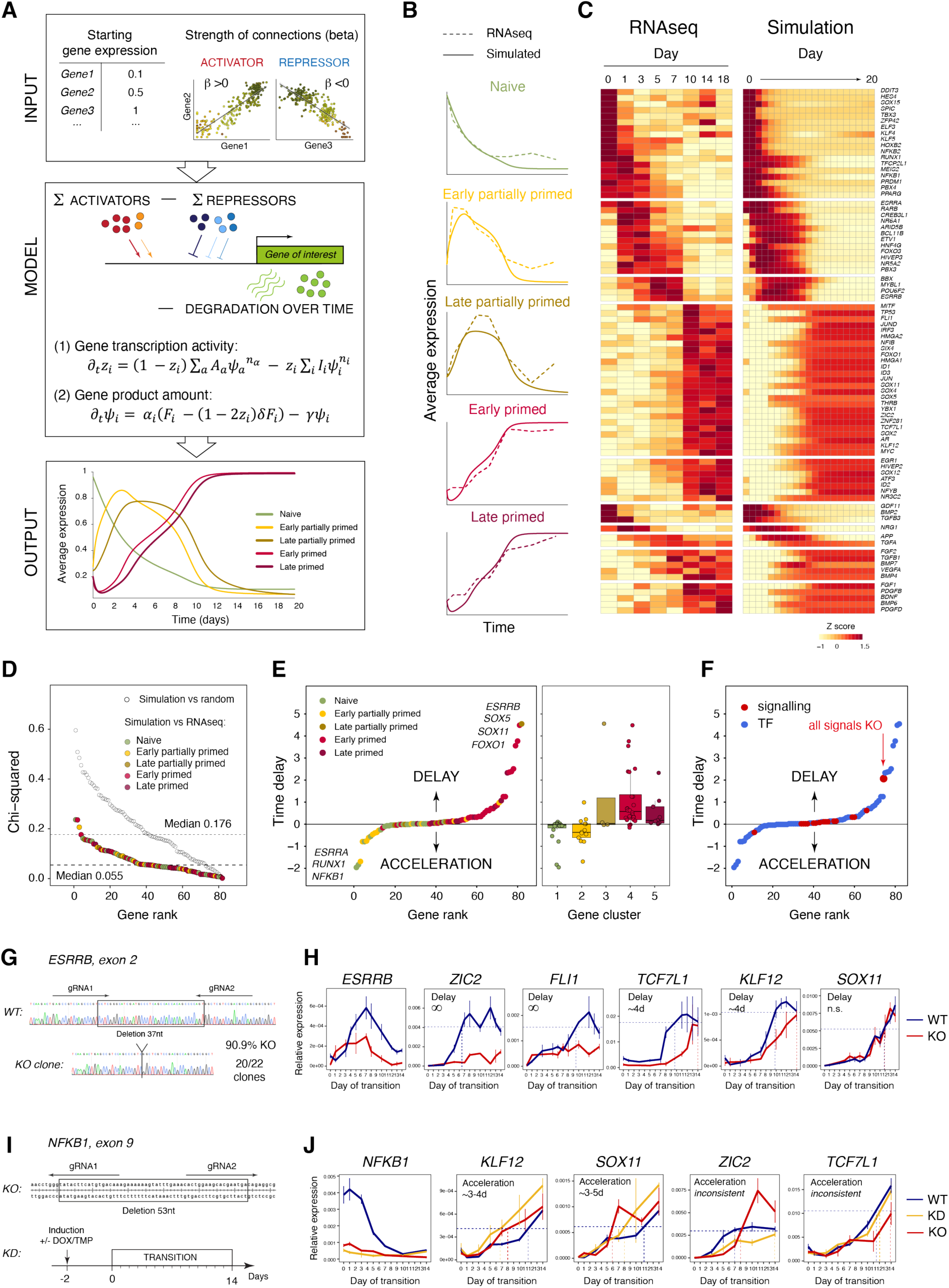
Transcriptional clock directionally guides the pluripotent state transition with defined timing (also see Figure S6 and Table S6). (A) Mathematical model of the pluripotent state transition. The result of simulation is shown as the dynamics of average activity of gene clusters. *z_i_* – transcription rate; *ψ_i_* – concentration of the gene product; *A_a_* and *ψ_a_* – strength of the regulatory connections and concentrations of activators; *I_i_* and *ψ_i_* – strength of the regulatory connections and concentrations of repressors; *F_i_* – production rate; ψ – degradation rate. (B) Comparison of gene cluster expression dynamics from the mathematical simulation with those observed in the scRNAseq data. (C) Comparison of individual gene expression dynamics from the mathematical simulation with those observed in the scRNAseq data. (D) Chi-squared test comparing the simulated gene expression dynamics with those observed in the scRNAseq for individual genes. Chi-squared values for random network were calculated as controls. (E) Changes in timing of the transition timing in simulations of single-gene knockouts; each dot represents an *in silico* knockout of an individual GRN member. (F) Simulation of knockouts for signalling factors, including an “all-signal” knockout, during the pluripotent state transition. (G) Strategy of CRISPR-based *ESRRB* gene knockout. (H) Dynamics of *ESRRB* and primed marker expression during the transition in the *ESRRB* knockout and the respective control. (I) Strategy of CRISPR-based *NFKB1* gene knockout and knockdown. (J) Dynamics of *NFKB1* and primed marker expression during the transition in the cells with *NFKB1* knockout and knockdown (from two independent experiments), along with the respective control (uninduced cells). The dashed lines in (H) and (J) indicate 90% level of gene expression in the primed control cells calculated as the average of all time points starting from day 10.

The model operates in two steps. First, it calculates the transcription level of a target gene which is proportional to the sum of activities of all its activators minus the sum of activities of all its repressors (with protein activities linked to their concentrations). Second, the level of gene transcription is used to estimate the amount of gene product that is changing over time due to degradation. The mRNA and protein levels were assumed to be proportional due to the lack of information on post-transcriptional control for each gene. The model outputs gene expression dynamics over time, either for individual genes or their groups. The simulation yielded gene expression dynamics closely resembling the ones observed in RNAseq, achieving a median chi-squared value of 0.055 among the GRN members, considerably lower than that using a random network (Figure 4B-D, Table S6). Therefore, the predicted activities of the TFs and signals are sufficient to describe the temporal dynamics of the system.

To further explore how the GRN functions, we simulated genetic perturbations of its components using our mathematical model. In these *in silico* knockouts (KO) simulations, the activity of a given gene was set to zero throughout the transition.

Surprisingly, none of the *in silico* KO of the GRN members blocked the pluripotent state transition (Figure 4E, Table S6), indicating the robustness and redundancy of the network. Instead, some perturbations altered the timing of the transition.

Interestingly, the removal of signalling factors from the network had only a minor effect on the timing of the transition, even when all signals were simultaneously switched off (Figure 4F). This suggests that TFs play a primary role in driving this autonomous network.

Among the *in silico* KOs, delays were more pronounced than accelerations, with a maximum delay of +4.55 days (*ESRRB*) and acceleration of -1.96 days (*NFKB1*). Three *in silico* KOs accelerated the transition by more than 1 day, all associated with the naïve or early partially primed TFs. Conversely, 11 perturbations delayed the switch to the primed state by over 1 day, all linked to KOs of the late partially primed or primed TFs.

To validate these predictions, we performed experimental perturbations of the top genes modulating the timing of the pluripotency transition and assessed its dynamics using qRT-PCR for marker genes. First, we generated naïve hPSCs with a KO of *ESRRB*, which was predicted to cause the largest delay (+4.55 days). Consistent with our predictions, key primed genes were upregulated with delays of over 4 days (*KLF12* and *TCF7L1*) or failed to reach the control level during the measurements (*ZIC2* and *FLI1*) (Figure 4G and 4H).

Next, we perturbed *NFKB1*, a naïve marker whose *in silico* KO was predicted to most strongly accelerate the transition. We used CRISPR-based KO and inducible knock-down (KD) approaches. The KD was efficiently induced during the transition, and the gene expression dynamics in the KO and KD was compared to uninduced cells. Some key primed genes (*KLF12* and *SOX11*) were upregulated more rapidly in the cells with reduced *NFKB1* as compared to the control, supporting our predictions (Figure 4I and 4J).

Additionally, we tested hPSCs with a KO of *TBX3*, predicted to have a negligible effect on the timing (-0.01 days). Matching our simulations, the dynamics of gene expression in the KO closely followed the control cells during the pluripotency transition (Figure S6).

In summary, our mathematical modelling and experimental approaches revealed that the network of TFs directionally guides the pluripotent state transition, providing time control, thus acting as an intrinsic transcriptional clock of pluripotency.

### The regulatory logic of the transcriptional clock of pluripotency

To elucidate the principles governing cell state transitions during epiblast development, we examined the structure of the transcriptional clock of pluripotency using our mathematical model. We aimed to identify the most important connections within the GRN and derive an optimised minimal network. While this network is less suited for detailed gene-level predictions due to reduced redundancy, it distils the core regulatory logic, offering a coarse-grained view of interactions between gene modules, essential for understanding the underlying mechanism.

An automated machine learning (ML)-based fitting procedure was employed to optimise the parameters for both whole-cluster and individual gene interactions in the mathematical model, revealing the most impactful connections to gene expression dynamics. Simulation of the pluripotent state transition using this ML-optimised model achieved enhanced alignment between the observed and simulated dynamics, with a median chi-squared value of 0.011 among the GRN members – an improvement over the original network (Figure 5A-5C, Table S6).

**Figure 5.**
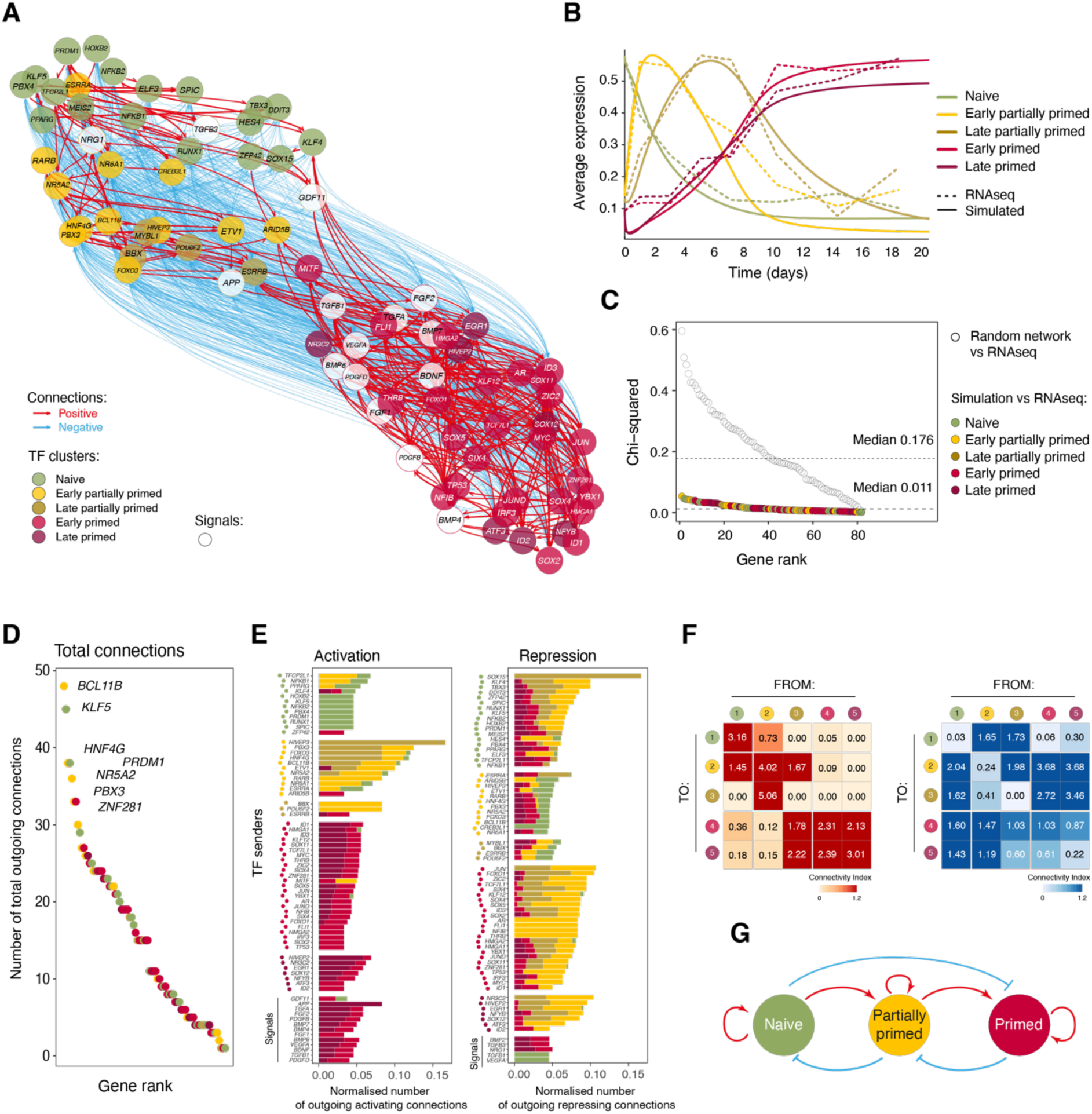
The regulatory logic of the transcriptional clock (also see Table S6). (A) UMAP representation of the ML-optimised GRN operating during the pluripotent state transition. (B) Comparison of gene cluster expression dynamics from the mathematical simulation using the ML-optimised model with those observed in the scRNAseq data. (C) Chi-squared test comparing the simulated gene expression dynamics using the ML-optimised model with those observed in the scRNAseq for individual genes. Chi-squared values for random network were calculated as controls. (D) Dot plot showing the numbers of regulatory connections between the ML optimised GRN members. (E) Bar plots showing the numbers of activating and repressing connections normalised to the total number of their connections, for each GRN member. The bar colors indicate the proportions of targets expressed at different stages of the transition. (F) Heatmaps showing the normalised numbers of activating and repressing connections between cell clusters. (G) The pluripotent state transition is a bistable regulatory system.

To investigate the regulatory logic driving the transcriptional clock, we analysed the interactions within our optimised GRN. This GRN contained 1,269 regulatory links, including 374 activating and 895 repressing, sent by 66 TFs and 16 signalling factors (between 1 and 48 outgoing connections per gene, median 15) (Figure 5D). We examined the distribution of the activating and repressing connections across stages, normalised to the total number of connections per cluster (Figure 5E and 5F). This revealed temporally short-range positive regulatory connections from earlier to later consecutive stages, as well as positive feedback within all clusters – except the late partially primed one, potentially due to the small size of this cluster. Furthermore, the network included short-range negative feedback loops and long-range repressive connections.

Overall, this analysis revealed a minimal interpretable coarse-grained model of the pluripotent state transition (Figure 5G). The regulatory logic of this transition resembled a bistable switch, where two exclusive states (naive and primed) self-reinforce and mutually repress one another. The naïve-to-primed transition is facilitated by a transcriptional cascade via a partially primed intermediate. This cascade relies on the short-range positive feedforward connections and negative feedback loops, enabling unidirectional and irreversible progression through defined stages: once the next step is ignited, the previous network is shut down. Additionally, the long-range repressive connections maintain the distinction between these molecular states, while the positive self-reinforcing connections stabilise them.

This regulatory logic explains the switch-like behaviour of the molecular network observed in our multiomics measurements. Importantly, the autonomous switch operates at a specific time point, consistent with our finding that this regulatory cascade provides temporal control for the transition thus acting as a transcriptional clock of pluripotency.

### Restructuring of the pluripotency clock explains differences in timing of epiblast development

A striking example of the differences in the timing of the pluripotent state transition is provided by a cross-species analysis: the epiblast development takes about 2 weeks in humans while only 2 days in mice^2–4^. We used this cross-species comparison to elucidate the mechanisms governing developmental timing.

We compared the transcriptional dynamics during the pluripotent state transition in hPSCs^21^ and mESC^47–49^ using published RNAseq datasets. Principal component analysis (PCA) effectively separated the species along PC1 (accounting for 70.6% of the total variation) (Figure 6A). Among the top genes contributing to PC1 were several factors known for their differing dynamics in human and mouse epiblast, including *KLF2*, *NROB1*, *TCF7L1*, *ESRRB*^18,32^. PC2 revealed the trajectory of the naïve-to-primed transition in both species (explaining 7.9% of the variation), with all three mouse datasets aligned in a correct temporal order along this trajectory.

**Figure 6.**
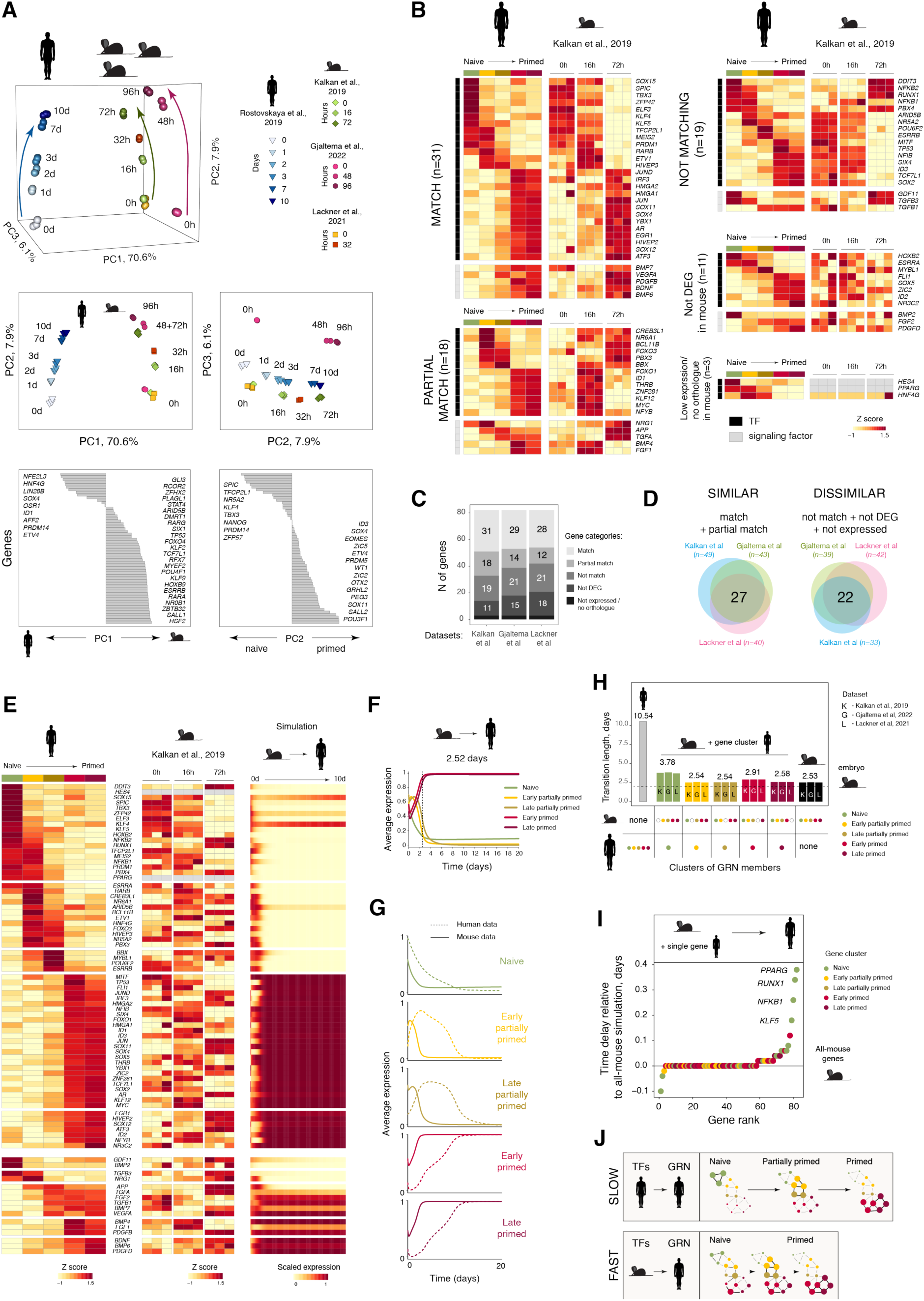
Restructuring of the pluripotency gene regulatory network explains the differences in timing of the epiblast development (also see Figure S7 and Table S7). (A) Principal component analysis of the trajectories of the pluripotent state transition in hPSCs and mESCs. (B) Expression of human pluripotency GRN members during the pluripotent state transition in mouse ESC (from Kalkan et al., 2019^47^), categorised based on their similarity to human gene expression dynamics. (C) Numbers of genes in each category of similarity to human gene expression dynamics, from three published RNAseq datasets on mouse pluripotent state transition. (D) Overlaps of genes within similarity and dissimilarity categories derived from comparisons of three published RNAseq datasets on mouse pluripotent state transition versus the human transition. (E) Expression dynamics of pluripotency GRN members in hPSCs and mESCs observed in RNAseq data, along with the simulated dynamics of the human pluripotency GRN using the mouse expression profile as input. (F) Gene cluster expression dynamics from the mathematical simulation of the human pluripotency GRN using the mouse expression profile as input. (G) Comparison of gene cluster expression dynamics from mathematical simulation of the human pluripotency GRN using either human or mouse expression profiles as input. (H) Timing of the pluripotent state transition from mathematical simulations of the human GRN where all-but-one gene clusters were set to mouse values, compared to simulations using all-human and all-mouse gene expression inputs. The simulations used three published RNAseq datasets from mESC transitions, with average timing indicated. (I) Dot plots showing differences in transition timing in simulations of the human GRN using all-but-one mouse gene expression input. Each dot represents the insertion of a human value for an individual GRN member into the mouse expression profile. The time difference was calculated relative to the all-human-genes transition. (J) Schematics showing the result: the GRN model derived for humans predicted the acceleration of developmental timing in mice when initialised with mouse-specific expression patterns, yet still leading to a human-like expression state.

Among the top contributors to PC2 were known naïve and primed pluripotency markers, including the members of our inferred GRN *TFCP2L1*, *KLF4*, *SOX11*, *ZIC2*^18^. Therefore, the general trajectory of the pluripotent state transition is conserved between mouse and human.

We compared the expression dynamics of the members of the pluripotency clock between the species (Figure 6B-6D, S7A and S7B, Table S7). Surprisingly, only 31.4% to 37.8% of the genes exhibited matching dynamics, that is changing in the same direction throughout the transition. An additional 14.6% to 21.9% were partially matching, i.e. changing in the same direction at least during a part of the process.

Conversely, the remaining 40.2% to 51.2% of the genes changed in the opposite direction, were not differential or not detected in mice. This observation is consistent with our previous analysis of human and mouse embryos (Figure S1), which showed that TFs are expressed in a different order in these species. This reordering indicates that the TFs exert their regulatory roles, both activating and repressing, at different stages of the pluripotency transition, which equates to a restructuring of the network. Therefore, the difference in timing between species is associated with the restructuring of the regulatory network rather than its simple scaling.

To investigate whether this restructuring can explain the differences in timing, we applied our mathematical model. We used transcriptome data from 3 independent mouse datasets^47–49^ as the initial expression values. These were combined with the regulatory relationships established in the human GRN, which serve as a proxy for TF interactions with specific cis-regulatory elements and regulatory connections of signal, reflecting the human GRN architecture. All other parameters were kept from the human model.

The simulated transition accelerated from 10.56 days with the human expression data, to 5.74-5.78 days with the mouse ones (Figure S7C), therefore the TF levels alone can partially explain the difference in developmental timing. Next, we increased the protein degradation rate in the model by a scaling factor of 2.3 to match the mouse rate, as previously reported^12^. Strikingly, the timing of the transition closely aligned with that of mouse epiblast development and mESC transition *in vitro*, lasting for 2.52-2.54 days (Figure 6E-6G, S7D and S7E, Table S7). Thus, “time-keeping” and “pace-making” mechanisms collectively explain species-specific developmental timing.

Additionally, we performed a series of simulations using various combinations of human and mouse gene expression levels as model inputs (Figure 6H). This revealed that the cluster of naïve markers contributed most significantly to the inter-species differences in timing. Specifically, genes such as *PPARG*, *NFKB1*, *RUNX1* and *KLF5* had the largest impact on accelerating the transition (Figure 6I and S7F, Table S7). Hence, the lower level of naïve genes in mice can explain the faster pace of the epiblast development and the *in vitro* pluripotency transition.

Furthermore, although the timing was similar to that of mice in these inter-species simulations, the final molecular profile matched the human characteristics (Figure 6J and 7). Therefore, while the levels of the GRN members account for developmental timing, the regulatory connections within the GRN (or its architecture) determine the final cell state.

**Figure 7.**
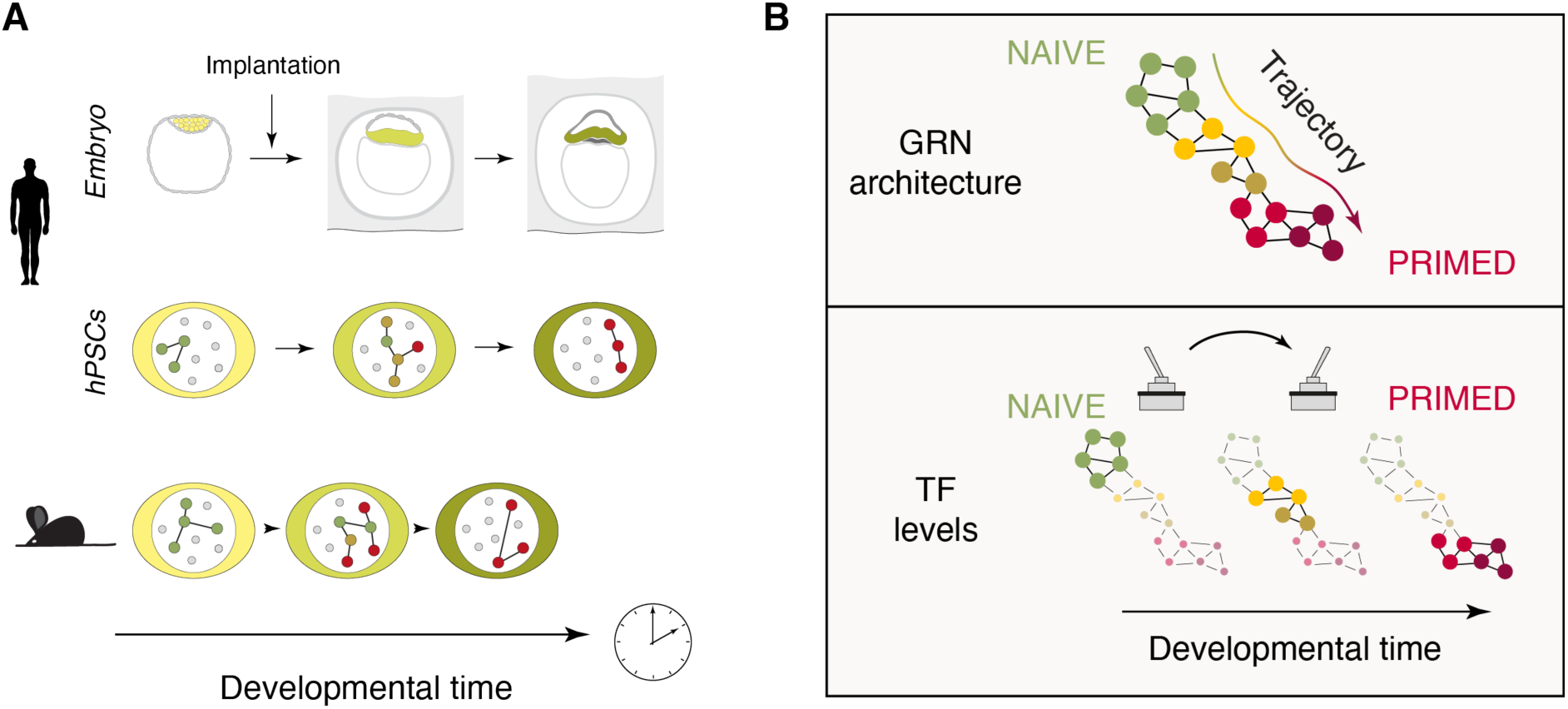
A transcriptional clock of the human pluripotency transition. (A) An intrinsic network of TFs drives the pluripotent state transition with defined timing, serving as a transcriptional clock. In mice, these TFs appear in a different order compared to humans, and the faster transition is associated with a restructuring of the GRN. (B) The pluripotency GRN directionally guides the pluripotent state transition, operating as a bistable switch. While the GRN architecture dictates the trajectory and final state, TF levels define timing.

To summarise, we demonstrated that the pluripotency transition is a robust, autonomous molecular programme encompassing a temporally defined cell state switch. By reconstructing the dynamic integrative GRN of TFs and signals combined with mathematical modelling, we identified a regulatory system governed by temporally short-range and long-range regulatory connections that follows the logic of a bistable switch. While the expression levels of the GRN members account for the developmental timing, the architecture of the regulatory connections defines the final molecular state. Therefore, this regulatory network not only directionally guides the developmental progression, but also acts as a transcriptional clock of pluripotency, intrinsically measuring time during the cell state transition.

## DISCUSSION

Timing is a fundamental concept in developmental biology, with important implications for understanding developmental mechanisms and embryonic malformations. In this study, we leveraged advanced GRN inference on multimodal single-cell data to reconstruct the molecular network of TFs guiding the human pluripotency transition. Through mathematical modelling, we created a precise digital twin of this transition, capturing its dynamics at a fine-grained level and enabling accurate predictions. Using this model, cross-species comparisons, and functional experiments, we demonstrated that this regulatory system schedules the cell state transitions and provides temporal control, acting as an intrinsic timer, which we have termed the transcriptional clock of pluripotency.

Surprisingly, we have found that the entire pluripotent state progression relies on an autonomous molecular programme. This was unexpected, given the extensive potential for self-organisation in the human embryo^50,51^ and in stem-cell-derived embryo-like structures^52–55^, which ultimately depend on intercellular cross-talk.

Nevertheless, among the three cell types that compose the blastocyst – the epiblast, primitive endoderm, and trophectoderm – the epiblast is arguably the most evolutionarily ancient and conserved. All developing vertebrate embryos pass through a transient pluripotency stage, including fish, frogs, birds^56–59^, whereas the trophectoderm is specific to mammals^60^. Moreover, the key players of the pluripotency network are highly conserved. For example, *Nanog*, *Pou5f3* (the homolog of mammalian *POU5F1*) and *SoxB1* (related to *SOX2*) are essential for developmental progression in the zebrafish pre-gastrulation embryo^61,62^. Therefore, it is likely that the molecular programme of the epiblast evolved earlier and independently of the other blastocyst cell types, maintaining its autonomy. However, we also showed that signalling factors do target the TFs of the pluripotency GRN, albeit they play a lesser role in its progression. We propose that signalling, including auto- and paracrine interactions, as well as intercellular cross-talk with other cell types in the embryo, helps fine-tune the pace of progression and coordinate development in the complex embryonic environment.

Our mathematical model and functional experiments demonstrated that the dynamic pluripotency GRN functions as an intrinsic timer during epiblast development (Figure 7A). Developmental timers can generally be categorised into two groups: oscillatory and linear rate-limiting (or “hourglass”) mechanisms. Oscillatory mechanisms, such as the somite clock^9^, circadian clock^63^, and cell cycle^64^, are driven by reactions with delayed negative feedback loops, resulting in periodic pattern. Hourglass mechanisms, including the transcriptional clock of pluripotency, rather rely on a threshold behaviour, where a factor triggers a downstream event once it reaches a critical level. For example, the mid-blastula transition and zygotic genome activation in *Xenopus* and *Drosophila* occur after several rounds of zygotic divisions, when the nuclear-cytoplasmic ratio reaches a specific threshold, measured through the concentration of certain maternal factors^5–7^. Similarly, the timing of *C. elegans* larval progression is determined by the concentration of miRNA *lin4* and its target *lin14*^8^.

Our mathematical simulations and experimental data suggest that *ESRRB* is a key component of the network, with its threshold level defining the time of the switch to the primed pluripotency. Consistent with our model’s predictions, hPSCs lacking *ESRRB* exhibited a substantial delay during the transition. However, the transition was not entirely blocked, indicating redundancy within the network and the existence of compensatory pathways. We hypothesize that this redundancy ensures the robustness of early development, which relies exclusively on this autonomous programme.

Our analysis of the GRN structure revealed that it follows a bistable switch regulatory logic. Bistable systems consist of two exclusive, self-reinforcing, and mutually inhibiting states. This regulatory logic underlies irreversible, directional developmental progression, as seen in neural progression^65^; and robust lineage choices, such as between the inner cell mass and trophectoderm^66^, or between mesoderm and neural fates^67^. Our findings suggest that it accounts for the distinction between the naive and the primed states, as well as their stability. Due to these regulatory connections, the epiblast progression follows a contingent developmental pathway and is irreversible. Therefore, the reverse, primed-to-naïve, conversion requires reprogramming, either *in vitro* or through the germline.

Transcriptional cascades have been proposed to act as developmental timers in various systems. One of the best-studied examples is the neuroblast clock in Drosophila^68^. In this system, neuroblasts undergo several cell divisions, producing four types of neurons in a temporally ordered, stereotypical sequence. The temporal information is conveyed by an autonomous cascade of TFs that rely on cross-regulation: *Hunchback* – *Krueppel* – *POU-domain protein 1* – *Castor*^69^. The specific type of neuron produced by the neuroblast depends on the TF expressed at the time of division. Manipulating Hunchback and Krueppel levels altered the timing of this developmental progression in neuroblasts^69–71^. Therefore, transcriptional cascades acting as intrinsic timers are likely common mechanisms across organisms.

Through mathematical modelling, we demonstrated that the levels of TFs, particularly at the naïve stage, account for the cross-species differences in the timing of epiblast progression (Figure 7B). Strikingly, a combination of “time-keeping” and “pace-making” mechanisms was sufficient to explain the developmental timing. While it remains to be determined how the naïve epiblast is equipped with a different set of TFs in different species, the differences in the dynamic GRN prior to the epiblast stage likely account for this. Furthermore, we showed that the order of TF expression differs between species; additionally, the TF regulatory connections directionally guide hPSCs to the human-like primed state. Mechanistically, these connections reflect interactions between TFs and their target cis-regulatory elements. Therefore, the architecture of the GRN is likely responsible for the trajectory and order of events during this developmental window. A key unanswered question is what consequences this heterochronic change poses for embryo development. One striking difference between mouse and human development is the order of lineage segregation from the epiblast. For example, the amnion and extraembryonic mesoderm segregate earlier in the human embryo as compared to the mouse^4^. We have previously shown that the competence to generate the early amnion exists transiently between the naive and primed states, matching the timing of its emergence in the embryo^34^. Therefore, it is possible that the transcriptional timer opens windows of competence, thereby explaining the differences not only in the timing, but also in the order of lineage segregation between species. This aligns with the role of other developmental timers in regulating responsiveness to signals and cell competence, such as the somite and neuroblast clocks.

Since developmental timers were first discovered in 1984^8^, several developmental processes have been found to rely on intrinsic molecular timekeeping mechanisms. Identifying the molecular mechanism that govern timing is critical for understanding fundamental processes and developmental defects, as well as for improving *in vitro* differentiation protocols, yet it remains a major challenge. It is still unknown whether time-keeping mechanisms operate throughout the entirety of development and across different lineages. Another intriguing question is whether post-natal processes, including ageing, are governed by molecular clocks and if interventions can be identified to alter their progression.

## LIMITATIONS OF THE STUDY

*In vitro* models may not fully recapitulate embryonic development.

The number of human embryos sequenced in Xiang et al., 2020^30^, was limited, and *in vitro* cultured embryos used for single-cell sequencing may differ from those developing *in utero*. Interspecies analyses are limited by existing knowledge of gene orthology.

Due to the high cost of 10X Multiome sequencing, only one replicate was performed. Although batch correction was applied, some batch effects cannot be entirely excluded due to their coincidence with biological differences. To mitigate these limitations, we validated our findings using independent sequencing datasets and gene expression assays.

Single-cell sequencing has limited sensitivity, which may result in inability to detect lowly expressed genes or less open chromatin regions.

The *in silico* ChIP relies on TF binding motif annotation, while the signalling pathway analysis is based on published evidence of regulatory connections. GRN inference relies on correlations of gene expression. Due to these limitations, false-positive or - negative connections between TFs and signals are possible, but validation is not feasible due to the lack of functional data. Our mathematical model and experimental validations support the overall robustness of the predicted regulatory logic and findings.

Our mathematical approach models the overall rate of gene activation and repression, summarising all regulatory molecular processes (e.g. epigenetic, post-transcriptional control). While this method does not resolve specific regulatory mechanism, it effectively captures the fundamental principles and gene expression dynamics.

## RESOURCE AVAILABILITY

### Lead contact

Requests for further information and resources should be directed to and will be fulfilled by the lead contact Maria Rostovskaya

### Materials availability

All unique/stable reagents generated in this study are available from upon request the lead contact with a completed materials transfer agreement.

### Data and code availability

The 10X Multiome data will be publicly available as of the date of publication. All original code will be available upon request.

Any additional information required to reanalyze the data reported in this paper is available from the lead contact upon request.

## ACKNOWLEDGEMENTS

The work was supported by the Babraham Institute’s BBSRC Core Capability Grant (BB/CCG2210/1), Institute Development Grant (BB/IDG2210/1) and Institute Strategic Programme Grant (BBS/E/B/000C0522); the MRC UK (MR/V02969X/1 to M.R. and W.R.); the FWO (“Fonds Wetenschappelijk Onderzoek”, fund for scientific research Flanders, VN423422N to L.C.); the ERC (EU Horizon 2020, no. 950349 to S.R.) and the Wellcome (210754/B/18/Z).

We wish to acknowledge Ricard Argelaguet for help and advice with *in silico* ChIP and GRN inference. We are grateful to Gavin Kelsey for resources and support. We thank Andrew Bassett, Sarah Cooper and Stuart Horswell (Sanger Institute, UK) for the advice and help with gRNA design. We thank Irene Zorzan for sharing protocols and reagents, Stephen Clark for technical advice, and Tom Owens for technical help. We thank the Babraham Institute Genomics Facility, Amelia Edwards, Connor Roberts and Paula Kokko-Gonzales, for help with library preparation; the CRUK Genomics Facility and Katarzyna Kania for sequencing; the Babraham Institute Bioinformatics Facility and Laura Biggins for initial data processing; Jeroen Galle from Ghent University for data handling and submission to the repository; the Babraham Institute Flow Cytometry Facility and its members for technical assistance. CRISPRi plasmids were a gift from Azim Surani, PX458 from Feng Zhang (both through Addgene); while pCMV-hyPBase from Allan Bradley.

## AUTHOR CONTRIBUTIONS

Conceptualisation, M.R., W.R.; Methodology, L.C., M.C., M.R.; Validation, A.K., M.R.; Formal analysis, L.C., M.C., M.R.; Investigation, L.C., M.C., A.K., M.R.; Data Curation, L.C., M.C., M.R.; Writing – Original Draft, M.R.; Writing – Review & Editing, L.C., M.C., T.M., S.R., W.R.; Visualisation, L.C., M.C., M.R.; Supervision, M.R., W.R., S.R., T.M.; Project Administration, M.R.; Funding Acquisition, M.R., W.R., S.R., T.M., L.C., M.C.

## INCLUSION AND DIVERSITY

One or more of the authors of this paper self-identifies as a gender minority in their field of research. We support inclusive, diverse, and equitable conduct of research.

---

**Figure S1.**
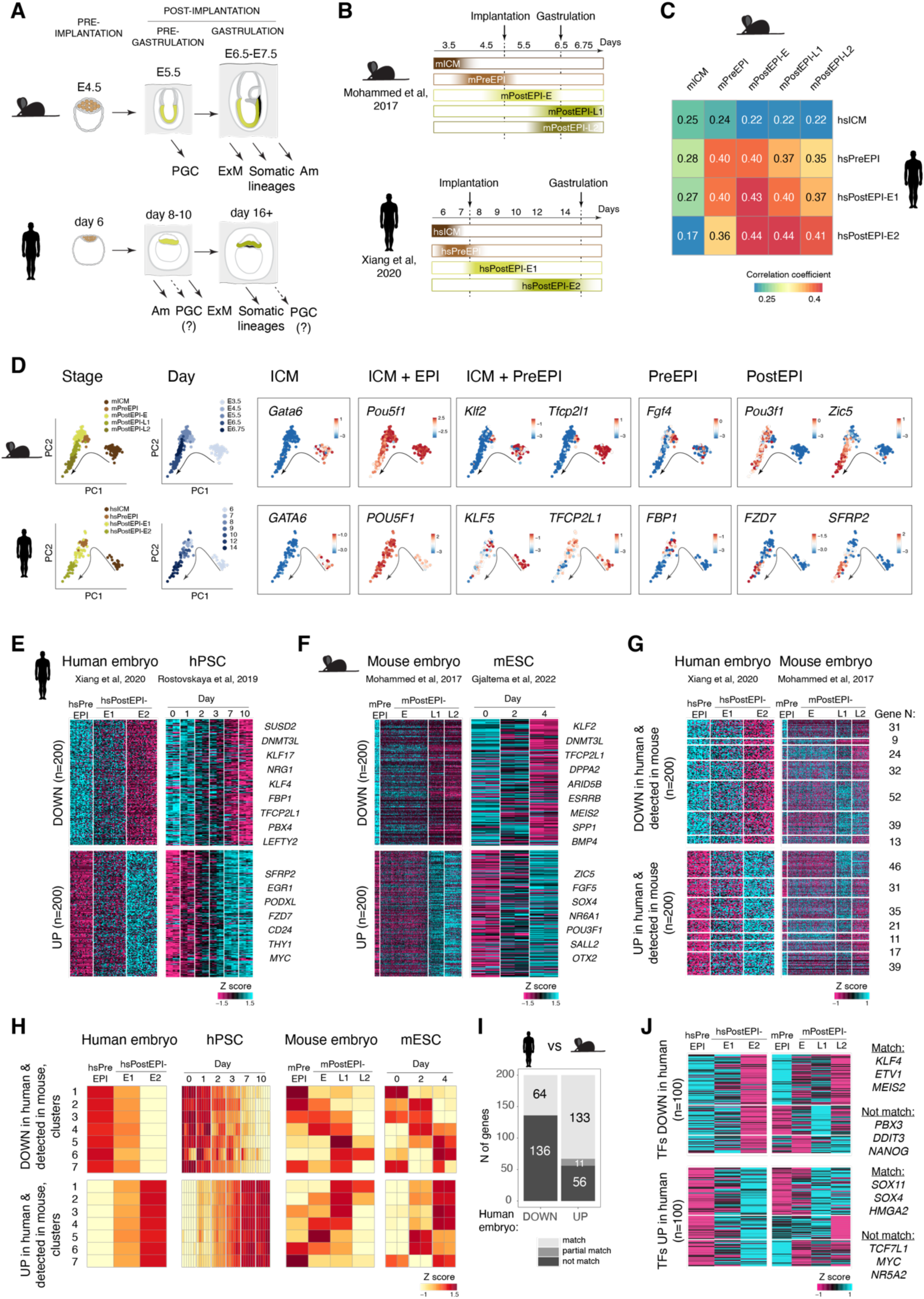
Pluripotent state transition in human and mouse embryonic epiblast (also see Table S1). (A) Schematics showing the timing and order of events in embryo development. (B) Representation of human and mouse epiblast stages from published single-cell RNAseq datasets (Xiang et al., 2020^30^; Mohammed et al., 2017^29^) used in this study. (C) Pearson correlation coefficients between human and mouse epiblast stages, computed from single-cell RNAseq. (D) Marker gene expression across epiblast development in human and mouse embryos. (E) Expression of the human epiblast top variable genes in hPSCs (published dataset from Rostovskaya et al., 2019^21^). (F) Expression of the mouse epiblast top variable genes in mESCs (published dataset from Gjaltema et al., 2022^49^). (G) Expression of the human epiblast top variable genes in mouse epiblast; filtered for expression in both species and clustered according to their expression dynamics in mouse. (H) Average gene expression levels in the clusters identified in (G). (I) Comparison of the gene expression dynamics in human vs mouse epiblast. Downregulated genes: clusters 1-3 matching, clusters 4-7 not matching. Upregulated genes: clusters 1-4 matching, cluster 5 partially matching, clusters 6-7 not matching. (J) Expression of the human epiblast top variable transcription factors in mouse epiblast (analysed as in (G)). Abbreviations: Am – amnion, PGC – primordial germ cells, ExM – extraembryonic mesoderm, ICM – inner cell mass, PreEPI – preimplantation epiblast, PostEPI – postimplantation epiblast, UP – upregulated genes, DOWN – downregulated genes.

**Figure S2 (related to Figure 1).**
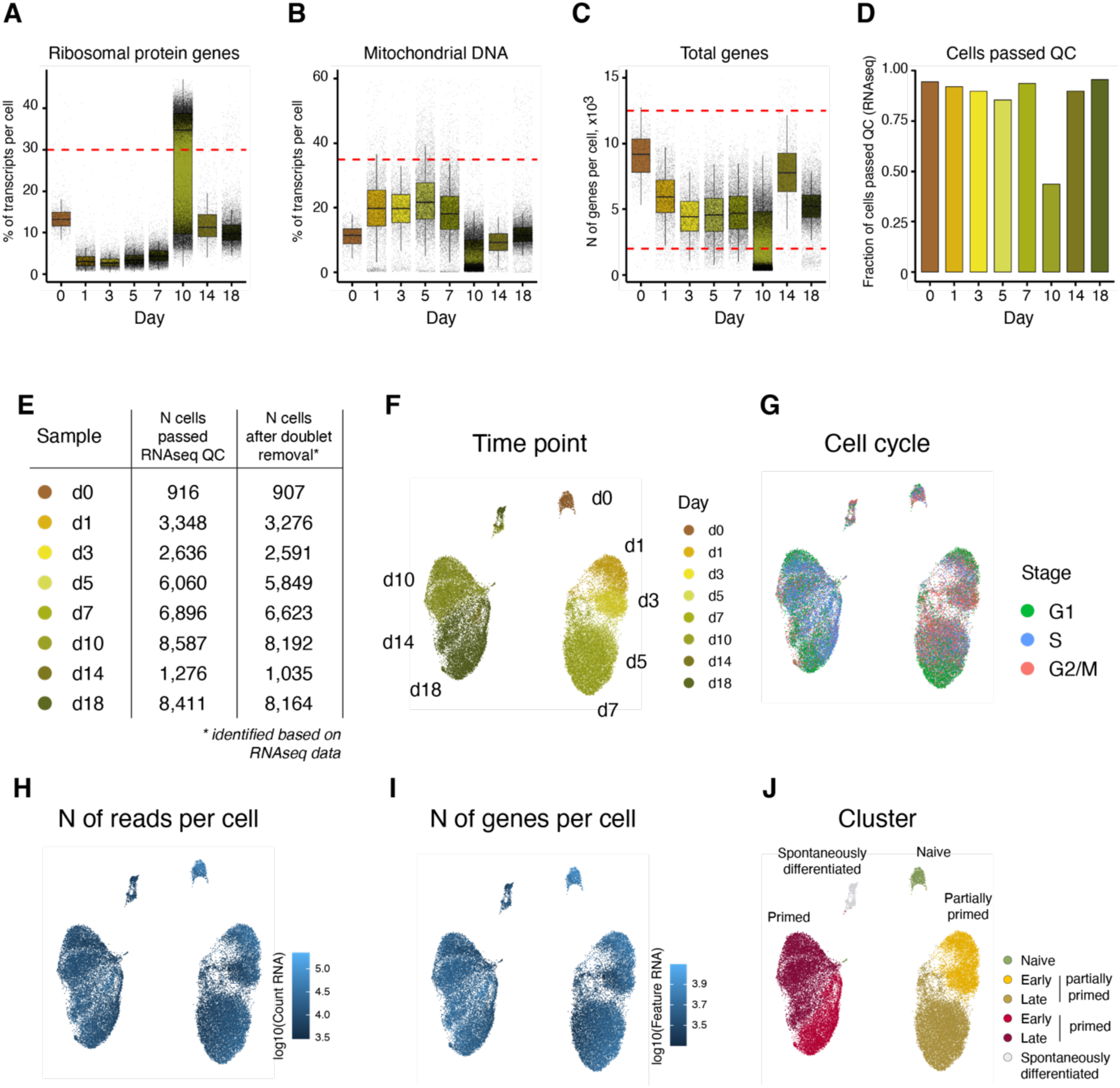
RNA expression profiling using 10X Multiome. (A) Box plot showing the percentage of ribosomal protein gene transcripts in each sample. (B) Box plot showing the percentage of mitochondrial RNA in each sample. (C) Box plot showing the number of genes detected per cell in RNAseq data. (D) Bar plot showing the fraction of cells passing quality control (QC) in RNAseq. (E) Table showing the number of cells passing QC and the number remaining after doublet removal for each sample in RNAseq. (F) UMAP based on RNAseq, with colors indicating transition days. (G) UMAP based on RNAseq, with colors indicating cell cycle stages. (H) UMAP based on RNAseq, with colors indicating read count per cell. (I) UMAP based on RNAseq, with colors indicating the number of genes detected per cell. (J) UMAP based on RNAseq, with colors indicating cell cluster.

**Figure S3 (related to Figure 1).**
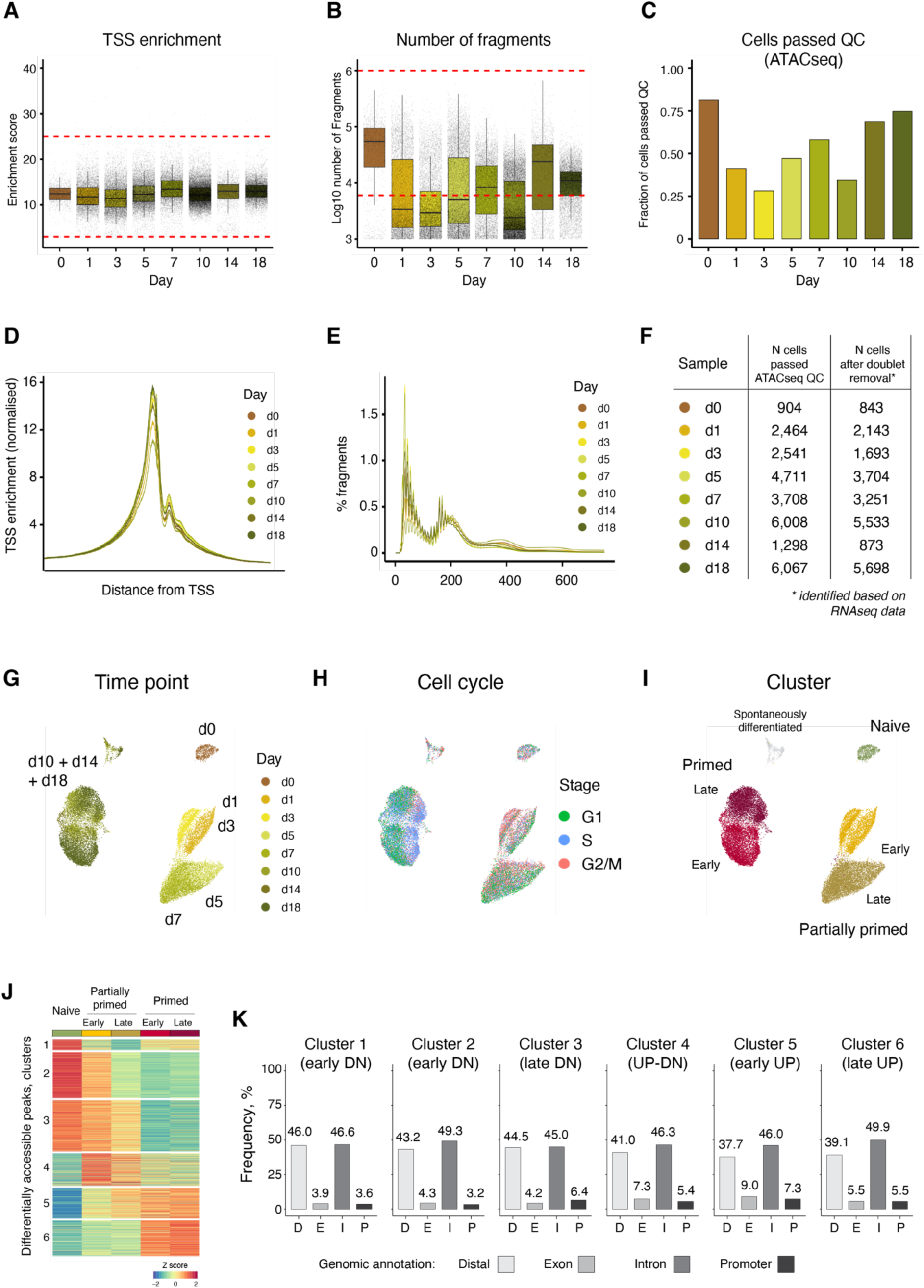
Chromatin accessibility profiling using 10X Multiome. (A) Box plot showing TSS enrichment per cell in ATACseq samples. (B) Box plot showing the number of fragments per cell in ATACseq samples. (C) Bar plot showing the fraction of cells passing QC in ATACseq samples. (D) Line plot showing fragment enrichment around TSS for each ATACseq sample. (E) Line plot showing fragment size distribution in ATACseq samples. (F) Table showing the number of cells passing QC and remaining after doublet removal for each sample in ATACseq. (G) UMAP based on ATACseq, with colors indicating transition days. (H) UMAP based on ATACseq, with colors indicating cell cycle stages. (I) UMAP based on ATACseq, with colors indicating cell clusters. (J) Heatmap from differential peak analysis using the ATACseq data. (K) Bar plot annotating peak clusters to genomic regions.

**Figure S4 (related to Figure 3).**
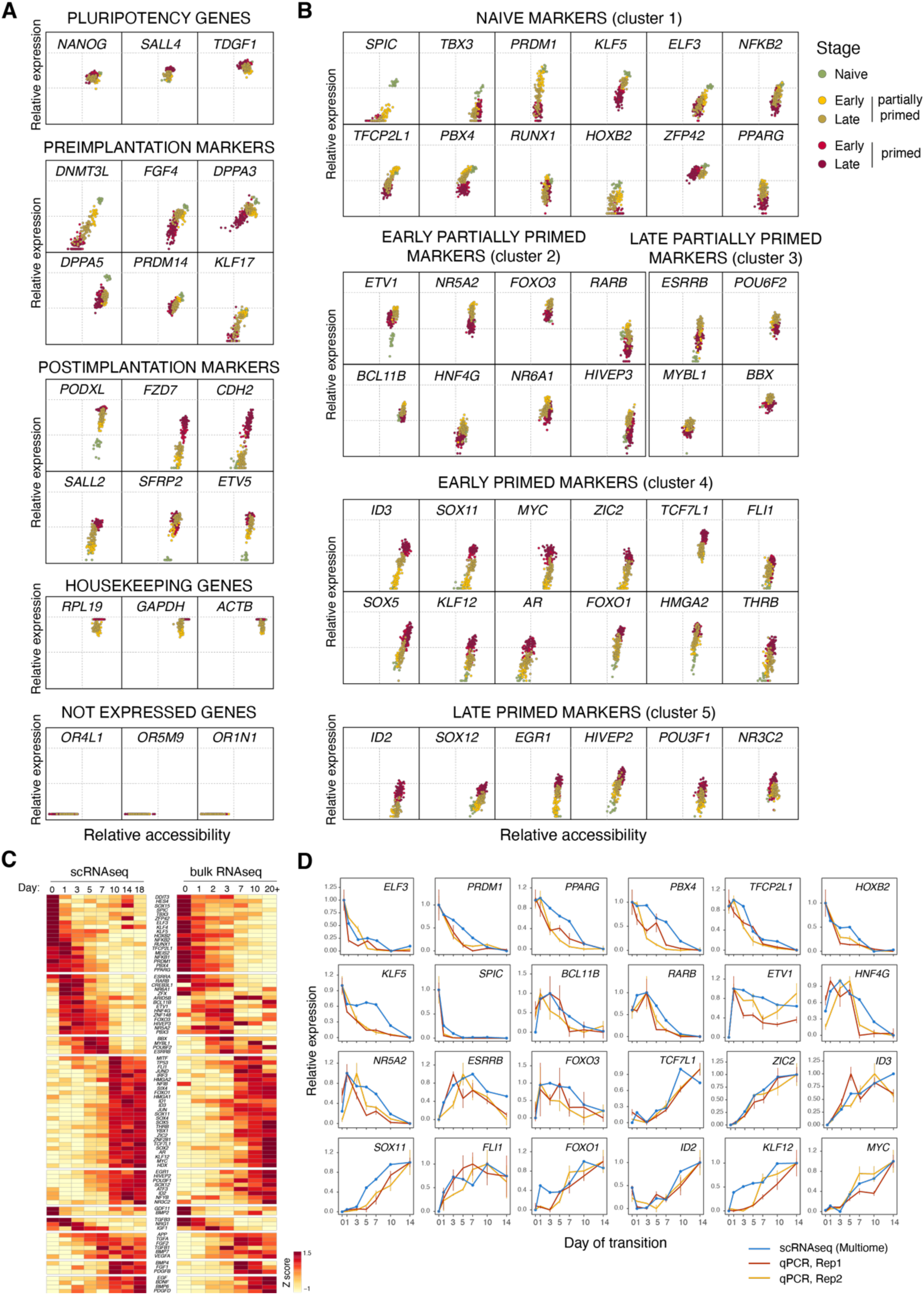
A catalogue of the most dynamically expressed TFs during the pluripotent state transition. (A) Plots showing the relative expression and accessibility of the pluripotency markers in metacells during the pluripotent state transition. Housekeeping genes were used as positive controls, olfactory receptors as negative controls. (B) Plots showing the relative expression and accessibility of selected TFs dynamically expressed during the pluripotent state transition. (C) Heatmaps showing the expression of the most dynamic TFs in the scRNAseq component of the 10X Multiome data in the present study and in published bulk RNAseq data (Rostovskaya et al, 2019^21^) (D) Validation of the expression profiles of dynamically expressed TFs as shown by qRT-PCR in two independent experiments.

**Figure S5 (related to Figure 3).**
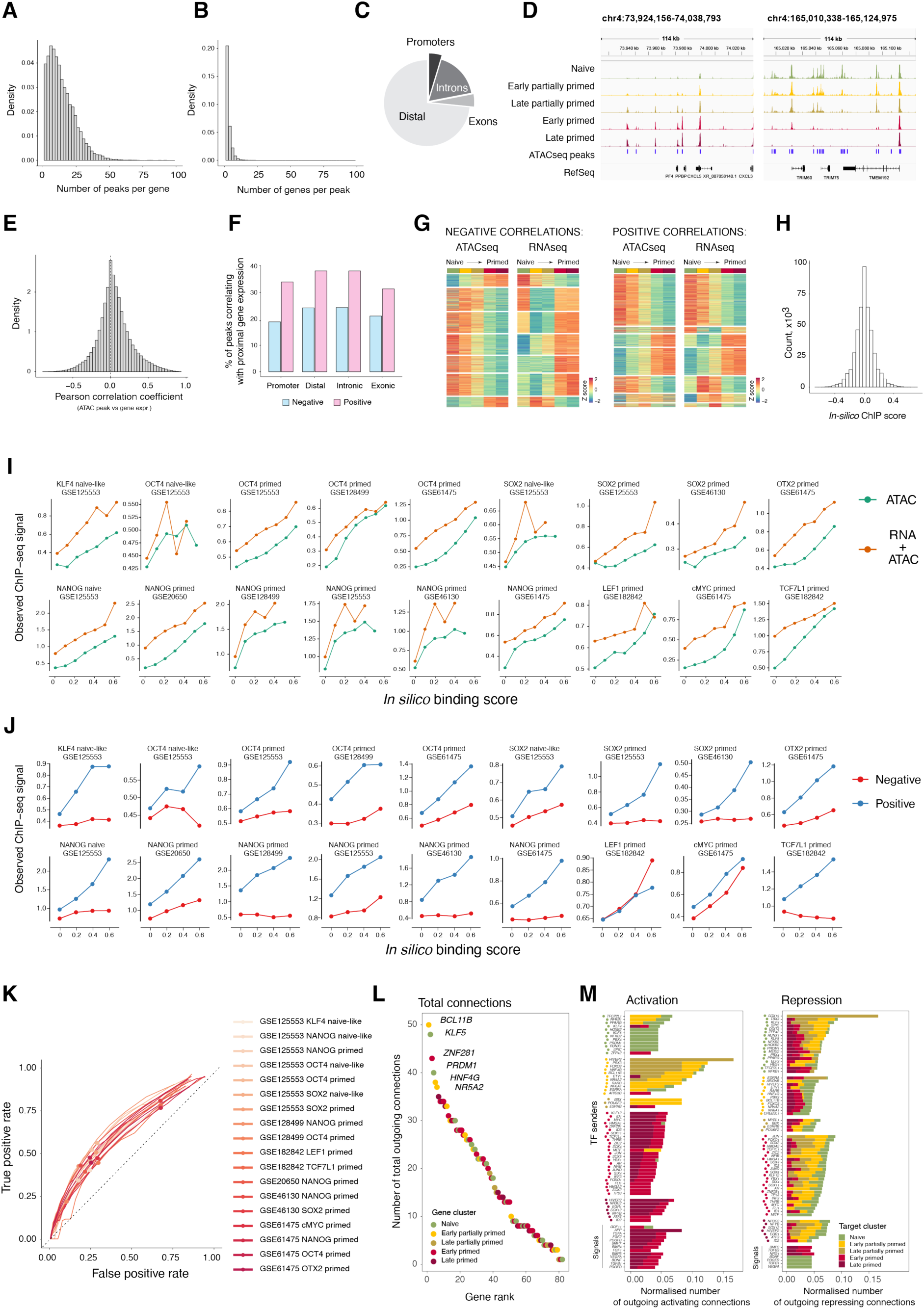
Construction of the *in silico* ChIPseq library for gene regulatory network inference. (A) Histogram showing the number of ATACseq peaks associated with each gene. (B) Histogram showing the number of genes associated with each ATACseq peak. (C) Genomic annotation of the ATACseq peaks associated with genes (located within 50kb). (D) Genome browser tracks for genomic regions with dynamically accessible peaks. (E) Histogram showing the Pearson correlation coefficient between ATACseq peak accessibility and the expression level of the associated gene. (F) Bar plot showing the proportions of ATACseq peaks per genomic category, whose accessibility positively or negatively correlates with the expression of the associated gene. (G) Heatmaps showing the ATACseq peak accessibility and the associated gene expression for pairs with positive and negative correlations. (H) Histogram showing the distribution of *in silico* ChIP binding scores. (I) Comparison of the *in silico* ChIP results obtained using RNAseq and ATACseq combined versus ATACseq data alone. The first strategy incorporates TF motif binding score, peak accessibility, and the correlation between TF expression and binding site accessibility, whereas the second omits this correlation. Cis-regulatory elements were predicted using each method at various *in silico* ChIPseq score thresholds, and the experimental ChIPseq signal was assessed within those regions. Regions identified using combined RNAseq and ATACseq data show a greater enrichment in the experimental ChIPseq data. (J) Experimental ChIPseq signal levels at binding sites predicted using *in silico* ChIP across various binding scores, shown separately for peaks with positive and negative *in silico* ChIP binding scores. Positive scores indicate a positive correlation between TF levels and peak accessibility, while negative scores indicate a negative one. Peaks with positive *in silico* binding score consistently exhibit greater enrichment of the experimental ChIPseq signal than those with negative scores. (K) Receiver Operating Characteristic (ROC) curves comparing TF binding sites predicted using *in silico* ChIP with those identified using published experimental ChIPseq data. Dots indicate an *in silico* binding score of 0.06. (L) Dot plot showing the numbers of regulatory connections between the GRN members. (M) Bar plots showing the numbers of activating and repressing connections normalised to the total number of their connections, for each GRN member. The bar colors indicate the proportions of targets expressed at different stages of the transition.

**Figure S6 (related to Figure 4).**
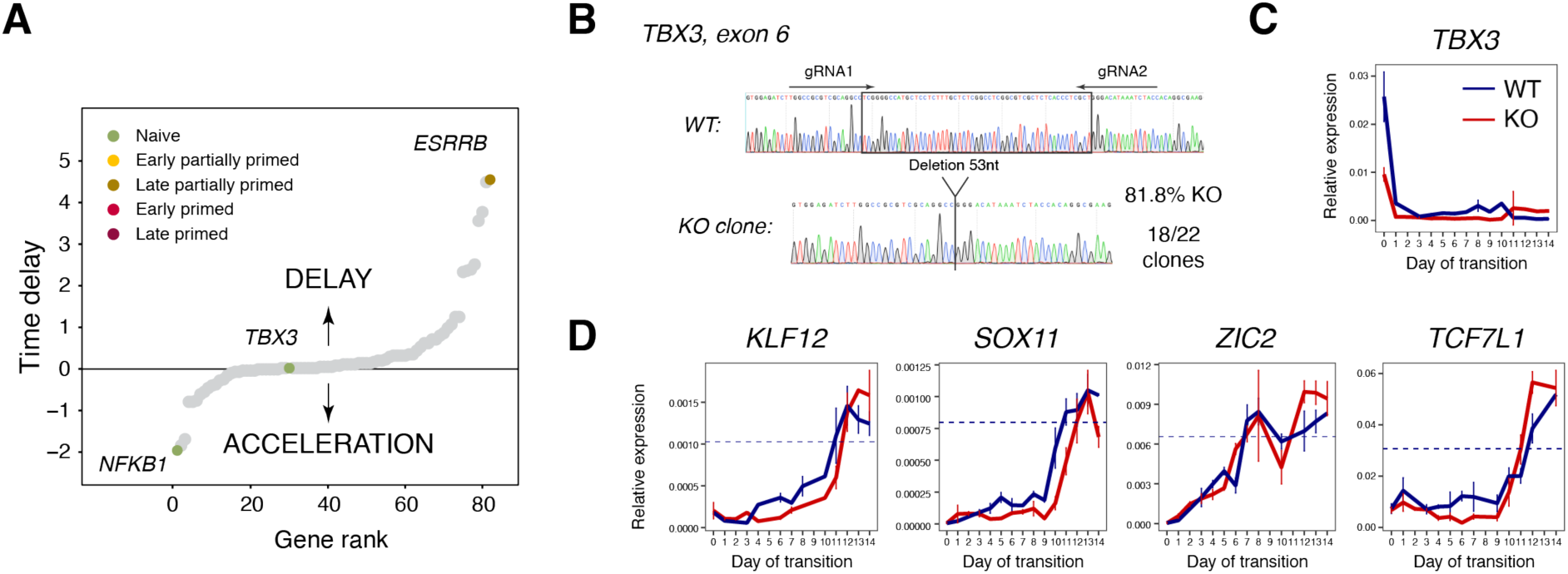
Mathematical model predicts TFs guiding the pluripotent state transition. (A) Changes in timing of the transition timing in simulations of single-gene knockouts; each dot represents an *in silico* knockout of an individual GRN member (as in Figure 4E). Top accelerating and delaying KO’s are shown, along with *TBX3*, predicted to be neutral. (B) Strategy of CRISPR-based *TBX3* gene knockout. (C) The level of TBX3 expression during the transition, in KO and the respective control. (D) Dynamics of gene expression in the *TBX3* knockout and the respective control during the transition. The dashed lines indicate 90% level of gene expression in the primed cells.

**Figure S7 (related to Figure 6).**
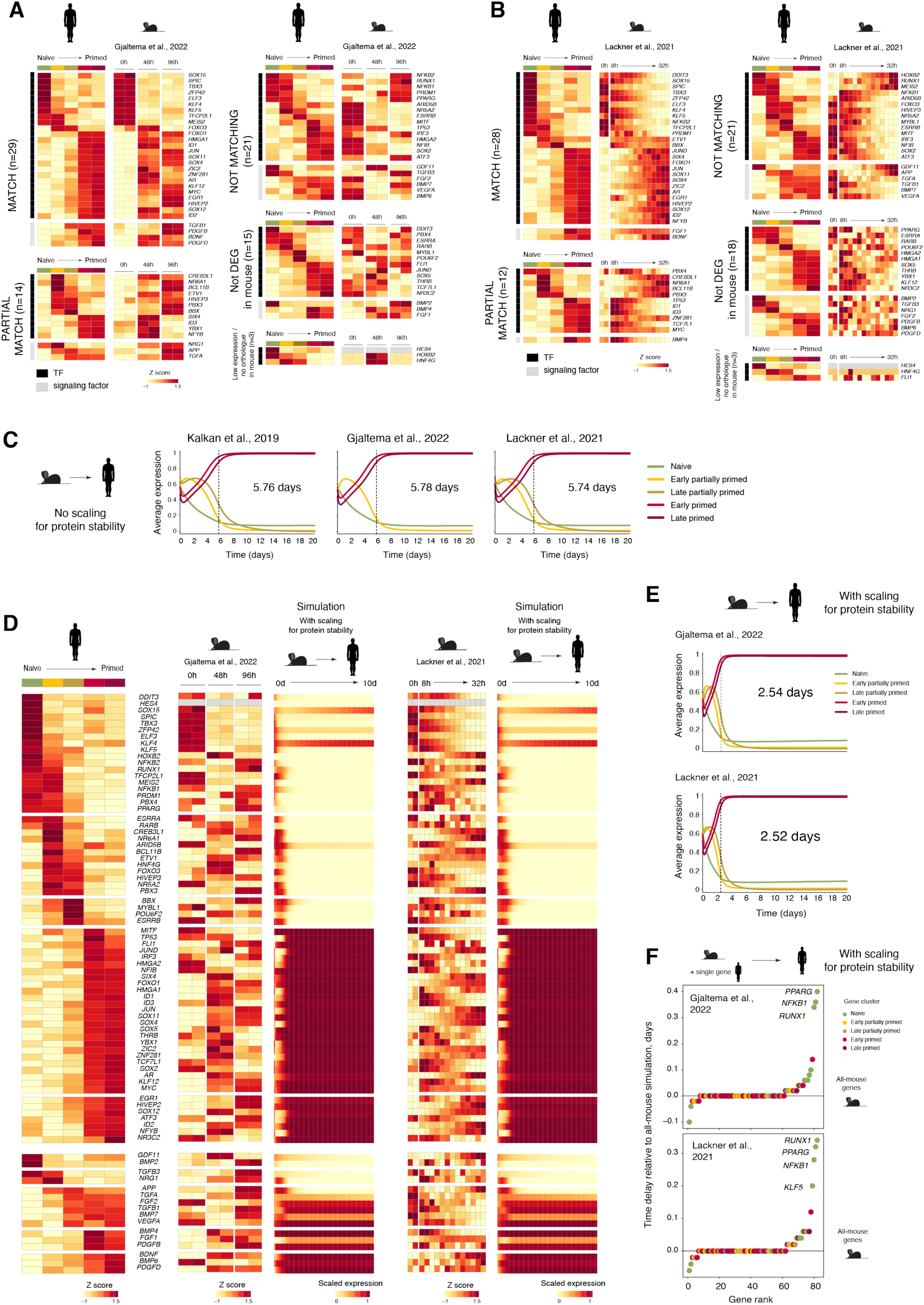
Restructuring of the pluripotency gene regulatory network explains the differences in timing of the epiblast development. (A) Expression of human pluripotency GRN members during the pluripotent state transition in mouse ESC (from Gjaltema et al., 2022^49^), categorised based on their similarity to human gene expression dynamics. (B) The same as (a), using the RNAseq dataset from Lackner et al., 2021^48^. (C) Gene cluster expression dynamics from the mathematical simulation of the human pluripotency GRN using the mouse expression profile as input and the human protein degradation rate. (D) Expression dynamics of pluripotency GRN members in hPSCs and mESCs observed in RNAseq data, along with the simulated dynamics of the human pluripotency GRN using the mouse expression profile as input and the scaled protein degradation rate (from Gjaltema et al., 2022^49^, and Lackner et al., 2021^48^). (E) Gene cluster expression dynamics from the mathematical simulation of the human pluripotency GRN using the mouse expression profile as input and the scaled protein degradation rate (from Gjaltema et al., 2022^49^, and Lackner et al., 2021^48^). (F) Dot plots showing differences in transition timing in simulations of the human GRN using all-but-one mouse gene expression input and the scaled protein degradation rate (from Gjaltema et al., 2022^49^, and Lackner et al., 2021^48^). Each dot represents the insertion of a human value for an individual GRN member into the mouse expression profile. The time difference was calculated relative to the all-human-genes transition.

## MATERIALS AND METHODS

### Cell lines

The study was conducted using the embryo-derived HNES1 (male) naïve hPSC lines^72^. The use of these hPSC lines was approved by the UKSCB Steering Committee. The research falls under Category 1A (“Exempt from review by a specialised oversight process” according to the ISSCR Guidelines for Stem Cell Research and Translation (2023).

### hPSC maintenance

Naïve hPSCs were cultured on irradiated mouse embryonic fibroblasts (MEFs) in PDLGX medium prepared as follows: N2B27 supplemented with 1μM PD032590, 10ng/ml human LIF (both from Cambridge Stem Cell Institute facility), 2μM Gö6983 (Tocris Bio-Techne, Cat. 2285), and 2μM XAV939 (Tocris Bio-Techne, Cat. 3748), as described previously^21,106^. The composition of the N2B27 basal medium was as follows: Neurobasal (Cat. 21103049, ThermoFisher Scientific) and DMEM/F12 (Cat. 31331093, ThermoFisher Scientific) in the ratio 1:1; 0.5% N2 (Cat. 17202048, ThermoFisher Scientific); 1% B27 (Cat. 17504044, ThermoFisher Scientific); 2mM L-glutamine (Cat. 25030024, ThermoFisher Scientific); 100mM 2-mercaptoethanol (Cat. M7522, Sigma-Aldrich). Geltrex (A1413302, ThermoFisher Scientific) was added at a concentration 0.5μl/ml to the culture medium during passaging.

Naïve hPSCs were passaged using TrypLE Express (Cat. 12604021, ThermoFisher Scientific) as single cells. 10μM ROCK inhibitor (Y-27632, Cat. 688000, Millipore) was added for 24 hours after re-plating.

All cells were cultured in a humidified incubator with 5% O_2_ and 5% CO_2_ at 37°C.

### Pluripotent state transition

The pluripotent state transition was performed as previously described^21,107^. Prior to the transition, naïve hPSCs were passaged once under feeder-free conditions to minimise the number of fibroblasts. For this, naïve hPSCs cultured on MEFs were dissociated with TrypLE Express and plated in the naïve hPSC maintenance medium supplemented with 10μM ROCK inhibitor on non-coated tissue culture grade plates. Geltrex was then added directly at a final concentration of 1μl/cm^2^. Once confluent, the cells were dissociated with TrypLE Express and seeded onto Geltrex-coated tissue culture plates at 1.6x10^4^/cm^2^ in the naïve hPSC maintenance medium with 10μM ROCK inhibitor. After 48 hours, the cells were washed with DMEM/F12 supplemented with 0.1% bovine serum albumin (BSA) (Cat. 15260037, ThermoFisher Scientific) and the transition medium (N2B27 supplemented with 2μM XAV939) was applied. The medium was refreshed every 1-2 days. The transition was carried out for 14 days unless stated otherwise.

The cells were passaged at confluency by dissociation using TrypLE Express and plating to Geltrex pre-coated dishes with a dilution 1:2. 10μM ROCK inhibitor was added for 24 hours after dissociation.

### Processing of published single-cell RNAseq datasets from human and mouse embryos

Embryo-derived RNAseq datasets were processed using Seurat package V4.0.1^83^.

Raw single-cell RNAseq data from *in vitro* cultured human embryos^30^ was obtained from GSE136447, considering only protein-coding genes for analysis. The cells were filtered based on the number of detected genes (≥4,500 genes per cell at log2FPKM > 2; the threshold chosen based on the overall gene detection distribution); and the proportion of the largest gene (< 5%). 523 out of 555 cells passed the quality control.

Single-cell RNAseq data from *in utero* mouse embryos^29^ was obtained from GSE100597, also considering only protein-coding genes. The cells were filtered following the original manuscript criteria: ≥4,000 detected genes and ≤10% of mapped reads assigned to mitochondrial genes, retaining 721 cells.

The cells from each dataset were clustered using a shared nearest neighbor (SNN) modularity optimization-based algorithm. For the human embryo data, clustering was performed using 23 principal components (PCs) with a resolution of 1; for the mouse embryo data, 7 PCs with a resolution of 0.5 were used.

The cell clusters were then assigned to cell types according to the known marker expression. The following cell types were considered for the human embryo data: hsICM (inner cell mass), hsPreEPI (preimplantation epiblast) and hsPostEPI-E1 and -E2 (two clusters of the post-implantation epiblast). For the mouse embryo, we considered: mICM (inner cell mass), mPreEPI (preimplantation epiblast), mPostEPI-E (early post-implantation epiblast), mPostEPI-L1 and -L2 (two clusters of the late post-implantation epiblast).

Pearson correlations between the human and the mouse stages were computed using the 500 most variable genes in each epiblast, considering their mean expression within each cell cluster.

### Comparison of gene expression dynamics between mouse and human epiblasts

We used single-cell RNAseq data from human and mouse embryos to compare gene expression dynamics between epiblasts of these species, applying one-to-one gene orthology based on gene symbols. Since single-cell RNAseq has limited power to detect lowly expressed genes, we considered genes that were detectable in both datasets. Detectable genes were defined as those expressed at log2(FPKM) > (-2) in at least 20% cells of at least one cell type. Based on this, 8,873 and 7,385 protein-coding genes were identified in the human and mouse datasets, respectively.

To characterise gene expression dynamics, we identified highly variable genes in human and mouse embryos using z-scores for the detectable genes across the epiblast stages. To compare the human epiblast expression dynamics to that of mice, we selected 200 top variable genes in the human epiblast that were also detected in mouse and examined their expression in the mouse epiblast. For this, hierarchical clustering of these genes was performed using a complete linkage method (hclust default settings within the pheatmap function). The resulting clusters were then categorised based on their similarity to human dynamics into three groups: matching (consistent dynamics throughout the time course), partially matching (consistent dynamics during part of the time course), not matching (opposite dynamics throughout the time course).

### Bulk RNAseq data from mESCs and hPSCs

Publicly available RNAseq datasets from mouse pluripotent state transition *in vitro* were obtained from GSE121148^47^, GSE145653^48^, and GSE167356^49^, and processed as described in^108^. In brief, alignment was performed using STAR (Spliced Transcripts Alignment to a Reference)^80^ against the GRCm39 release 110, followed by summarising gene-level counts using FeatureCounts. All count files were annotated using biomaRt with mouse genome informatics (MGI) gene symbols, then normalised to the library size as counts per million (cpm), and log-transformed.

For human pluripotent state transition *in vitro*, the data were processed as described previously^21^, the dataset is available in GEO:GSE123055. Reads were aligned to human genome build GRCh38/hg38 with STAR 2.5.2b^80^ using human gene annotation from Ensembl release 87^109^. Htseq-count^110^ was subsequently used to quantify expression to gene loci.

Principle component analysis (PCA) was done using Factoextra 1.0.7 package^92^, considering 1,000 variable genes.

### Nuclei preparation

Nuclei preparation was performed according to 10X Genomics protocols.

All reagents were purchased from Sigma-Aldrich, Merck, unless stated otherwise.

For nuclei preparation, hPSCs were dissociated into single cells using TrypLE Express. After washing with phosphate-buffered saline (PBS) containing 1% BSA, the cells were filtered using 40μm strainer, pelleted by centrifugation, resuspended in PBS with 0.04% BSA and counted.

To obtain nuclei, a total of 10^6^ cells were pelleted by centrifugation for 3 minutes at 300g and resuspended in 200μl of cell lysis buffer containing 0.1% NP40, added to the lysis dilution buffer (10mM Tris HCl pH 7.4; 10mM NaCl; 3mM MgCl_2_; 1% BSA; 1mM DTT; 1U/μl RNasin). Lysis was performed for exactly 1 minute on ice, followed by a washing step using 500μl of washing buffer (0.1% Tween-20 in the lysis dilution buffer). The nuclei were pelleted by centrifugation for 3 minutes at 500g at 4°C. 500μl of PBS with 1% BSA and 1U/μl RNasin were added to the pellet, followed by incubation for 5 minutes. The nuclei were pelleted again (3 minutes at 500g at 4°C) and resuspended in PBS with 1% BSA and 1U/μl RNasin. Single nuclei were flow sorted after staining with 10μg/ml 7-aminoactinomycin (7-AAD) (Molecular Probes, ThermoFisher Scientific). The obtained nuclei were washed with PBS with 1% BSA and 1U/μl RNasin, followed by permeabilisation. For this, the nuclei were treated with 50μl of permeabilisation buffer containing 0.01% NP40, 0.01% Tween-20, and 0.001% digitonin in the lysis dilution buffer for 2 minutes on ice.

Alternatively, cell lysis and nuclei permeabilization were performed in a single step. For this, dissociated hPSCs were flow-sorted to obtain a single-cell suspension, followed by two washing steps using PBS with 0.04% BSA. A total of 10^5^ cells were pelleted (3 minutes at 500g at 4°C) and treated with 45μl of cold lysis buffer containing 0.025% NP40, 0.025% Tween-20, 0.0025% digitonin, and 1U/μl RNasin in the lysis dilution buffer for 0.5-3min on ice.

The permeabilised nuclei were washed three times: first, using 200μl of washing buffer, followed by two washes with 45μl of Diluted Nuclei Buffer (PN-2000153, 10X Genomics). The resulted nuclei pellet was gently resuspended in 10-20μl of Diluted Nuclei Buffer. A total of 17,400 nuclei in 5μl was used for library preparation.

### 10X Multiome library preparation and sequencing

10x Genomics single cell library preparation and relevant quality control was carried out by the Babraham Institute Genomics Facility. Each sample contained 17,400 nuclei. The samples were processed into Gel Beads in Emulsion (GEMs) using the Chromium Controller, following the manufacturer’s guidelines. Based on the input nuclei concentration, the maximum number of nuclei was targeted per sample (10,200-10,440).

Following the GEM generation, the Multiome library preparation for RNAseq (whole transcriptome) and ATACseq (chromatin accessibility) libraries was completed, following the manufacturer’s instructions, using the following kits: Chromium Next GEM Single Cell Multiome ATAC + Gene Expression Reagent bundle (PN-1000283/285); Chromium Next GEM Chip J Single Cell Kit (PN-1000234/1000230); Single Index Kit N Set A (PN-1000212) and Dual Index Kit TT Set A (PN-1000215).

Library quality control (QC) steps were completed using an Agilent 2100 Bioanalyser with the High Sensitvity DNA Assay (Agilent, Santa Clara, USA) for the cDNA generation and final libraries. The final libraries were quantified by qPCR with the KAPA qPCR Library Quantification Kit (Roche, KAPA Biosystems, Cape Town, South Africa).

Sequencing was conducted by the CRUK Cambridge Institute. Libraries were pooled and sequenced following the manufacturer’s instructions for 10x Genomics libraries on the Illumina NovaSeq6000 platform using S1 or S2 flow cells (Illumina, San Diego, USA).

Sequencing specifications were as follows. Gene expression libraries were pooled and sequenced as read 1: 28 cycles, read 2: 90 cycles, index 1: 10 cycles and index 2: 10 cycles. The ATACseq libraries were pooled separately and sequenced as read 1: 50 cycles, read 2: 49 cycles, index 1: 8 cycles and index 2: 24 cycles.

### 10X Multiome data analysis

10X Multiome data processing and data analysis followed the general framework developed by Argelaguet et al.^38^ (https://github.com/rargelaguet/mouse_organogenesis_10x_multiome_publication), including RNAseq and ATACseq analyses, *in silico* ChIP, and the GRN inference, with project-specific modifications. Additional custom analyses were implemented *de novo*. The resulting pipeline is detailed below.

### Single-cell RNAseq data processing

Sequencing files from the pluripotent state transition, along with previously obtained data from naïve HNES1 cells (GSE178379^96^) were processed using CellRanger^82^ with default arguments, based on the GRCh38 genome (v. 2020-A, Ensembl98). To ensure quality data, cells were selected based on the following criteria: (i) expressing between 2,000 and 12,500 features, (ii) with less than 30% of mapped reads originating from ribosomal proteins (based on gene annotation, RPL & RPS proteins) and (iii) less than 35% of mapped reads with a mitochondrial origin. The resulting count matrix was stored as a SingleCellExperiment object^84^. Doublet detection was done using the scds package, based on a combination of a co-expression-based and binary classification approaches^87^.

### Dimensionality reduction and cell cluster identification using RNAseq data

Principal component analysis (PCA) calculating 15 principal components (based on elbow plot) for the RNAseq data on the 2,500 most variable expressed genes was performed for linear dimensionality reduction. Subsequently, we relied on the uniform manifold approximation and projection (UMAP) algorithm^111^ on calculated principal components for visualization considering 50 nearest neighbors and a distance parameter of 0.25. Finally, Louvain clustering was performed based on 15 nearest neighbors and with a resolution of 0.25.

### Single-cell ATAC sequencing data processing and peak calling

ATACseq fragment files from hPSCs during the pluripotent state transition, along with previously obtained data from naïve HNES1 cells (GSE178379^96^) were generated by CellRanger^82^. Subsequently, the ArchR package^88^ was used to create arrow files. Filtering was performed to retain quality cells using the following criteria: (i) between 6,000 and 1,000,000 fragments, (ii) a transcription start site (TSS) enrichment between 3 and 25 and (iii) a maximal blacklist ratio lower than 0.03. Next, doublet annotation based on RNA-seq data was used to filter possible doublets.

Peak calling was performed using MACS2^89^. A consensus peak set was obtained through an iterative peak-overlapping strategy using ArchR. Peaks were annotated using ArchR’s default approach with the hg38 genome assembly.

Chromatin accessibility dynamics was analysed at individual peaks, gene promoters and broad gene loci regions. Promoter accessibility was assessed by aggregating peaks from 500bp downstream to 100 bp upstream of the transcription start site (TSS) using the ArchR’s gene score functionality (with tile size = 100bp, gene scaling factor for gene length = 1).

Gene accessibility was analysed by aggregating peaks from the genic region (TSS to the transcription end site) and surrounding up- and downstream sequences as defined by the standard ArchR settings. In brief, a region from 5kb upstream of the TSS was considered for all genes, with an additional extension of 1-10kb up- and downstream depending on the genomic context defined for each gene. The contribution of these distal regions was downweighted using a decay function (e⁻ˣ/5000 + 1/e, where x is the distance to the TSS in nucleotides). As per the standard settings, a tile size of 500 and gene length scaling factor of 5 was applied.

### Dimensionality reduction and cell cluster identification using ATACseq data

Iterative latent semantic indexing (LSI) was performed for dimensionality reduction of ATACseq peaks, with a projection in 25 dimensions for the 15,000 most variably accessible peaks. Harmony batch correction^112^ was performed using the ArchR package. Leiden clustering (as is default for ArchR) was performed with a resolution of 0.15 to define clusters.

### Multi-modal dimensionality reduction and cell cluster identification

Cells that passed quality control for both modalities separately were retained for multi-modal analysis.

To obtain a multi-modal dimensionality reduction, first linear dimensionality reduction was performed for individually measured modalities: (i) a PCA for RNAseq data including 15 principal components on the 2,500 most variable genes and (ii) an LSI for ATACseq data including 30 dimensions on the 25,000 most variable peaks, with a harmony batch correction^112^. Next, we generated a multi-modal latent embedding using MOFA+ with a fixed set of 30 factors^35,90^.

The MOFA result was inputted to the uniform manifold approximation and projection (UMAP) algorithm^111^ to generate a non-linear two-dimensional visualization, and was used for hierarchical cell clustering.

### Differential expression and accessibility analysis

The sparse nature of single-cell data limits the possibilities for statistical analysis; therefore, we aggregated RNAseq read or ATACseq fragment counts to generate “pseudobulk” replicates. This pseudobulk approach is particularly recommended for ATACseq as counts for peaks are typically even sparser than for RNAseq data. This approach also addresses the issue of different sample number per group, facilitating differential analysis and avoiding false discoveries^113^.

We performed bootstrapping 30% of cells creating five pseudo-replicates per cluster^114^. The analysis was performed similarly for gene expression and for chromatin accessibility at both the peak and gene levels. After aggregating counts, normalisation and log transformation were performed using scran^85^ and scuttle^86^ to obtain log-transformed counts per million (cpm).

Differential analysis was performed using the quasi-likelihood F-test implemented in edgeR^91^. Results were considered significant differentially expressed with a false discovery rate (FDR) <0.01 and a log2 fold change (|logFC|) ≥1.

Hierarchical clustering of differentially expressed genes was performed using Rclusterpp package based on Euclidian distances between z-scores.

Gene set analysis and pathway analysis was performed via the R API for Webgestalt^115,116^ through overrepresentation analysis against background of all annotated genes.

### Metacell selection

The sparsity of single-cell data limits analysis of the molecular dynamics and interactions, yet pseudobulk approach described above does not retain the information about cell heterogeneity. To overcome these limitations, we identified groups of similar cells, or metacells, and aggregated the single-cell sequencing data from those groups.

Metacells were inferred using SEACells software^36^, with the number determined by a target of 100 cells per metacell. The algorithm converged after 21 iterations (convergence epsilon of 1e-4) yielding 316 metacells (1.3% of total cells) with a median of 78 cells per metacell.

After QC (at least 750,000 RNAseq reads and 600,000 ATACseq fragments per metacell), 250 metacells were retained for further analysis with a median of 85.5 single cells per metacell. Each metacell was assigned to a cell cluster based on its initial seed cell’s cluster label.

### Reconstruction of the trajectory of the pluripotent state transition

The trajectory of the pluripotent state transition was inferred based on the gene expression values in metacells.

For metacell trajectory inference, highly variable genes were selected based on a gene expression variance p-value of <0.01, calculated using scran package. Pseudotime was calculated as a linear combination of the first principal component from the principal component analysis (scater package, 25 components) and the first diffusion coefficient from the diffusion map (destiny package^94^). The diffusion coefficient was scaled using a factor of 100 to match the range of the principal component.

### Visualisation of gene expression and accessibility in metacells

For visualisation, gene expression and chromatin accessibility values were min-max scaled, considering genes with log-transformed expression between 0 and 10 and chromatin accessibility between 1 and 8, to minimise the impact of outliers. Visualisation was performed using heatmaps^93^ and scatter plots.

### Transcription factor motif annotation

DNA binding molecules and their motif annotations were extracted from CISBP^104^; TFs were selected from this list based on the AnimalTFDB database^105^.

Motif matches for each peak were obtained using motifmatchr (v1.16.0), with a minimum motif width of 7 and a maximum q-value of 1e-4.

### Selection of the most dynamically expressed transcription factors

Differential gene expression analysis was performed using a pseudobulk approach, as described above. TFs were considered as the most dynamically expressed if they were upregulated in a given cell cluster compared to at least three out of four other cell clusters. Next, we assigned these TFs to the corresponding cell clusters or stages of the transition. Due to our approach, the dynamically expressed TFs could be specific to one or two cell clusters. In the latter case, we assigned the TFs to the earlier stage of the transition. As such, we identified 5 groups of TFs corresponding to the 5 temporal stages of the pluripotent state transition: naïve, early and late partially primed, early and late primed.

### Signalling pathway dynamics analysis

A combination of existing tools was used to analyse the dynamics of signalling pathways.

First, we used CellPhoneDB to examine ligand-receptor interaction dynamics during the pluripotent state transition^39–41^. CellPhoneDB provides a curated database of ligand-receptor interactions. The gene expression data from single cells were used as an input for this analysis. Dynamic ligand-receptor interactions were identified through the enrichment of ligand-receptor co-expression within cell clusters compared to random, based on single-cell label permutations.

Next, we used NicheNetR^42^ to predict target genes of these dynamic ‘ligand-receptor’ pairs, considering differentially expressed genes as potential targets. NicheNetR assigns weights to ‘ligand-target’ and ‘receptor-target’ interactions based on supporting evidence, factoring in quality and recurrence. We selected ‘ligand-receptor-target’ combinations with a weight above 0.10.

We then performed a linear regression analysis for ligand-target and receptor-target pairs, based on their expression in metacells during the pluripotent state transition. We considered the regulatory potential of ‘ligand-receptor-target’ combinations significant if they met the following criteria: (1) the expression dynamics of a ligand and its putative target should correlate; (2) the receptor’s dynamics should not contradict that of the respective signal (i.e., the receptor should either change in the same direction or remain unchanged); using thresholds of beta-value >0.24 and p-value <0.1. The cut-offs were selected to match the respective values used for the GRN inference (see section “Gene regulatory network (GRN) inference”).

### Genome-wide cis-regulatory element inference using in silico ChIP

TF binding can be experimentally identified using antibody-based methods coupled with sequencing, such as chromatin immunoprecipitation sequencing (ChIPseq) or CUT&Tag. However, these datasets are available only for a limited number of TFs and do not cover all cell types. We applied a previously reported *in silico* ChIP method^38^ to infer TF binding, based on the simultaneous measurement of gene expression and chromatin accessibility in aggregated metacells.

We considered ATACseq peaks as potential regulatory elements. TF binding was predicted based on (1) the presence of the TF binding motif within the region (using motifmatchr as described above), and (2) the correlation between the chromatin accessibility of this region and the expression level of the TF. Each ‘TF-regulatory element’ combination was quantitatively assessed using an *in-silico* ChIP score, as following^38^:

χ_ij_ = σ_ij_ minmax(θ_ij_π_i_), where

χ_ij_ – *in silico* ChIP binding score;

σ_ij_ – the correlation between the chromatin accessibility of peak *i* and the expression level of the TF *j* across the metacells;

θ_ij_ – the motif binding score of the TF *j* in peak *i*;

π_I_ – the maximum chromatin accessibility of peak *j* across the metacells.

This resulted in a comprehensive *in-silico* ChIP library comprising 781 TFs, their 2,086,946 potential target cis-regulatory elements (with a median of 1,680 regions per TF) and 32,255 unique target genes (with a median of 4,912 genes per TF).

To validate the *in silico* ChIPseq approach, we compared the result with publicly available ChIPseq datasets obtained in naïve or primed hPSCs for a set of TFs including KLF4, NANOG, POU5F1, SOX2, LEF1, TCF7L1, cMYC, and OTX2. Raw fastq files were obtained via the sra toolkit, trimmed using Trim Galore (v. 0.6.7) and aligned to GRCh38 reference genome (release 32) using Bowtie2^79^. Peaks were called by MACS2 (using “--broad and --broad-cutff 0.1” arguments) without an input control. The result was processed using samtools to create bam files^117^ and bamCoverage to create Bigwig files^118^ for visualisation.

First, we compared the performance of the *in silico* ChIPseq based on two strategies: (1) based on RNAseq and ATACseq data, as described above, and (2) based on ATACseq data alone, which relies only on TF motif binding score and peak accessibility, while omitting the correlation between TF expression and accessibility of its binding site. Cis-regulatory elements were predicted using each method at various *in silico* ChIPseq score thresholds, and then experimental ChIPseq signal was assessed within those regions.

We focused on the *in silico* ChIP approach combining both data modalities due to its superior performance, and evaluated it across different threshold scores. Cis-regulatory elements were predicted for various positive and negative *in silico* ChIP thresholds, and the ChIPseq signal observed in experimental data was assessed within those regions.

Next, we constructed receiver operating characteristic (ROC) curves for those TFs where ChIPseq data were available, using the latter as the ground truth. For this, the true positive rate (TPR) and the false positive rate (FPR) were calculated for different *in silico* ChIPseq score values, in order to identify an *in silico* ChIPseq score with the best TPR/FPR ratio. An *in silico* ChIPseq score of 0.06 was selected as a threshold for putative binding sites, enabling identification of 50% of ChIPseq peaks at TPR/FPR ∼ 2.

### Gene regulatory network (GRN) inference

The integrative dynamic GRN was reconstructed by considering interactions between dynamically expressed TFs and signaling molecules predicted to regulate these TFs. To infer ‘TF-target’ pairs, we surveyed our *in-silico* ChIP library for putative regulatory regions bound by the TF within 50kb of the target gene body, with a score above 0.06. ‘Signal-target’ predictions were made as described in the “Signalling pathway dynamics analysis” section.

The strength of all putative regulatory connections was quantitatively assessed using linear regression, modelling target gene expression as a function of sender gene expression in metacells. This approach assumes linear relationships between interacting genes throughout the transition, which may not always hold, particularly given that the transition involves 3 distinct cell states. To overcome this limitation, we optimised a system to select a subset of metacells for linear regression analysis as described below.

We considered the following clusters of metacells: naïve, early partially primed, late partially primed; while early and late primed metacells were combined into a single category due to their high similarity. For each ‘sender-target’ combination we removed metacells where the sender and its target do not interact, as these are not relevant for assessment. To achieve this, we identified the clusters to which the sender and target were assigned and excluded all metacells from the clusters that appeared after the later of the two.

Additionally, we implemented specific rules to TF interactions to harness cell heterogeneity for their quantification. We reasoned that TFs interact locally within single cells, whereas signalling factors are diffusible and act on the entire cell population. Consequently, local cell heterogeneity may not be informative for intercellular ligands. First, for TFs that are assigned to the same metacell cluster, we considered only these metacells for linear regression. Second, if a target gene was expressed earlier than the sender, we included only the metacells spanning the transition between these two stages omitting both the earlier and the later ones, as these were less relevant for the assessment.

Subsets of metacells used for linear regression:

| Cell clusters |  | Sender TF |  |  |  |
| --- | --- | --- | --- | --- | --- |
|  |  | 1 | 2 | 3 | 4+5 |
| Target | 1 | 1 | 12 | 123 | 12345 |
|  | 2 | 12 | 2 | 23 | 2345 |
|  | 3 | 123 | 123 | 3 | 345 |
|  | 4+5 | 12345 | 12345 | 12345 | 45 |

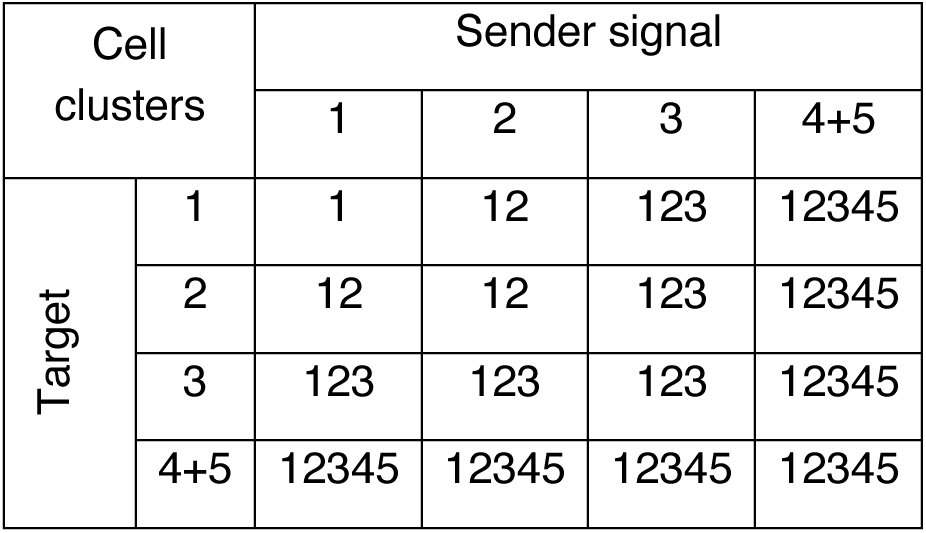

| Cell clusters |  | Sender signal |  |  |  |
| --- | --- | --- | --- | --- | --- |
|  |  | 1 | 2 | 3 | 4+5 |
| Target | 1 | 1 | 12 | 123 | 12345 |
|  | 2 | 12 | 12 | 123 | 12345 |
|  | 3 | 123 | 123 | 123 | 12345 |
|  | 4+5 | 12345 | 12345 | 12345 | 12345 |

Cell clusters: 1 – naïve, 2 – early partially primed, 3 – late partially primed, 4+5 – combined early and late primed.

As a result, we quantitatively assessed all predicted interactions withing the dynamic gene regulatory network governing the pluripotent state transition. Regulatory connections with positive beta-values were classified as activating, while those with negative beta-values were classified as repressing. To construct a regulatory network with contingent gene activation throughout the transition, only genes receiving at least one activating incoming interaction were retained, except for those of the initial naïve stage. Based on this, to retain the maximum number of genes in the network without compromising stringency, regulatory connections were selected using an absolute regression slope (beta value) above 0.24 and a p-value <0.1. The same thresholds were applied in the signalling pathway analysis (see section “Signalling pathway dynamics analysis” above).

For GRN visualisation, we performed PCA using 10 principal components, followed by UMAP transformation (spread = 0.5, minimal distance = 0.5, 30 nearest neighbors) on the TF expression matrix in metacells (modified from Fleck et al., 2023^119^). The resulting UMAP representation of the network was generated using ggraph package.

### Mathematical model of the pluripotency GRN

The mathematical model of the pluripotency GRN relied on the equations of motions (e.o.m.) for the gene activities, which were then used to simulate the network dynamics.

The master equation formulation for the network dynamics was based on gene switching between on and off states and protein production^120^. The network consists of *N* genes indexed by *i*. For each gene, we consider a binary degree of freedom *s_i_* ∈ {0,1} representing active and inactive states, and a degree of freedom *n_i_* ∈ ℕ describing the number of protein molecules produced.

Stochastic switching between the active and inactive states happens with rates *α_i_* And *β_i_*, respectively

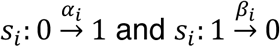

The rates *α_i_* and *β_i_* depend on the protein number *n_i_*. For describing gene expression simplifying assumptions were made for the model.

First, it was assumed that levels of mRNA molecules and proteins are proportional, given the lack of data on post-transcriptional control.

Secondly, to describe interactions between genes, we assumed that activators and repressors bind cis-regulatory elements at a protein-dependent rate, and unbind at a constant rate. The binding of activators was assumed to be both necessary and sufficient for gene activation, so that there is no spontaneous gene activation without activating transcription factors. That is, while binding of repressors leads to the inactivation of the target gene, their unbinding does not directly lead to inactivation.

Finally, we incorporated cooperative binding of proteins to DNA, and therefore a nonlinear dependence of the activation rates on protein abundancies.

Mathematically, for gene *i* regulated by protein *j*, the switching rates between active and inactive states are described as follows. The activating switching rates are given by:

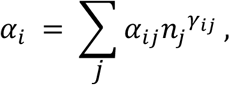

where *α_ij_* is the binding rate of activator protein *j* to the regulatory regions of gene *i*, and *γ_j_* is a degree of cooperativity of protein binding to DNA. This choice of rates corresponds to an activation by binding of a *γ_ij_*-mer of the regulating protein. Conversely, the rates of stochastic deactivation are given by:

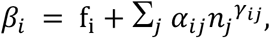

where *β_ij_* is the binding rate of repressor protein *j* to the regulatory regions of gene *i* at cooperativity *γ_ij_* and *f_i_* is the unbinding rate of activating proteins. Again, repression happens through the binding of a *γ_ij_*-mer of the repressing protein.

Protein production was defined by the following transition rates:

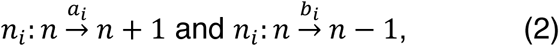

Where *a_i_* and *b_i_* represent protein production and degradation rates, respectively. The dependence of the production rate on the protein numbers was described in terms of *a_i_* = *F_i_* + *δF_i_* when gene *i* is active, and *a_i_* = *F_i_* − *δF_i_* when gene *i* is inactive, where *δF_i_* represents the offset in mRNA activation rate between these states and *F_i_* models the average protein production between active and inactive state. The degradation rate *b_i_* is independent of the gene state.

The state of the system is described by the probability of observing a protein configuration {*n_i_*} and gene activation state {*s_i_*} at a given time *t*, denoted by *P*({*n_i_*}, {*s_i_*}, *t*). This probability follows the master equation corresponding to the reaction rules defined in Eqs. (1) and (2),

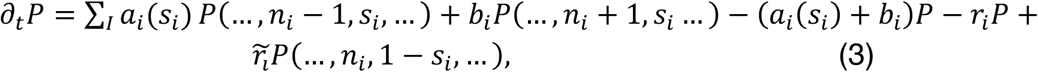

where *a_i_*(*s*) and *b_i_* are the protein production and degradation rates that depend on the gene state *s_i_* as defined above, and *r_i_, r̃_i_* indicate the rates of transition from the state *s_i_* to 1 − *s_i_* and vice versa.

The master equation (3) can be reformulated in an operator form by defining the single-gene state as:

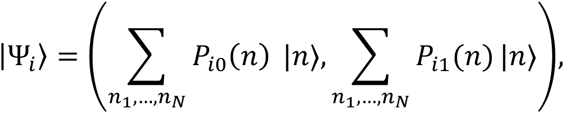

where *P_i_*_-_(*n*) and *P_i_*_1_(*n*) are the probabilities of observing the gene in the active or inactive state with n proteins, and expressing the right-hand side of (3) as the action of an operator on the system state describing the time evolution of the system through transition between different possible states as described by the introduced rates.

The total state of the system is given by:

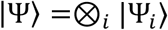

Where ⊗*_i_* indicates the tensor product between different single-gene state. This allows for a second-quantization formalism of the master equation Eq. (3), where the dynamics of the system state vector |Ψ⟩ follows:

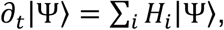

Where *H_i_* is the single-site evolution operator defined by:

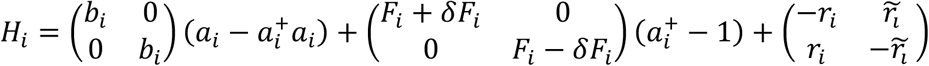

With *a_i_*|*n_i_*⟩ = *n_i_*|*n_i_* − 1⟩ and *a*^2^|*n_i_*⟩ = |*n_i_* + 1⟩.

The second quantization formalism enables the formulation of state transition in terms of an action functional^120^. The transition probability from an initial state |n*_i_*.⟩ at time 0 with protein numbers *n_i_*. to a final state |Ψ_3*i*._⟩ at time t with protein numbers *n*_3*i*._ reads:

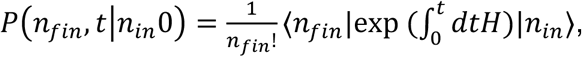

This can be reformulated in a path-integral formalism, where:

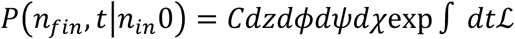

And the Lagrangian ℒ is given by:

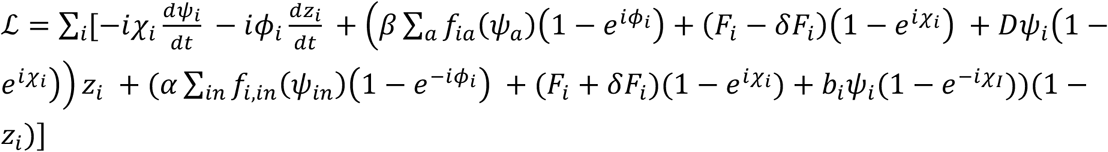

From this path integral, the equations of motion for average activities *z_i_* and protein number *ψ_i_* were obtained using a semiclassical approximation:

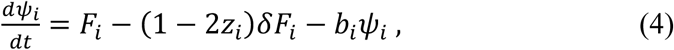

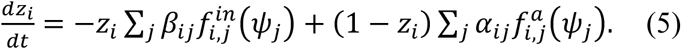

where *ψ_i_* is the average protein number for gene *i*; *z_i_* is the average fraction of time the gene *i* is active. 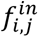 and 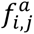 are the contributions of activators and repressors to the activity dynamics, respectively, and are in general of the form 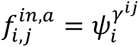 (coming from the cooperative binding of regulators introduced in the activation and repression rates), where the exponent *γ*^!$^represents the degree of cooperative binding of proteins to DNA.

In the following, we will set a gene-independent protein degradation rate *b_i_* = *b*, basal protein production rate *F_i_* = *F*, and an increase for activation *δF_i_* = *δF* for all *i*, and a gene-independent cooperation exponent *γ_ij_* = 2 for all *i*, *j* (as TFs frequently act as dimers). In this way, we aimed to reduce the number of parameters in the model without overly limiting the phenomenology of the model. This approximation is valid if degradation rates and cooperation exponents are similar across genes. Finally, time is measured in units of protein degradation rate, thus setting *b* = 1 and expressing all rates in these units.

In the approximation of fast protein dynamics with respect to the gene state dynamics, we can set the average protein number to its steady state, determined by 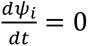, which gives:

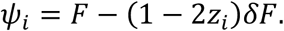

Considering this, we obtain the following equation for the time evolution of gene activities:

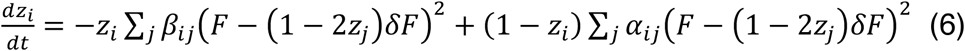

Therefore, this equation describes the gene activity dynamics as influenced by the presence of other regulating genes. The structure of Eq. (6) constrains gene activities to lie between 0 and 1.

### Simulation of the gene regulatory network dynamics

Data structures and numerical simulations were implemented using the Julia programming language^121^. The coefficients *α_ij_* and *β_ij_* in Eq. (6) are determined by the gene network structure, so that *α_ij_* ≠ 0 if gene *j* is an activator of gene *i*, and *β_ij_* ≠ 0 if gene *j* is a repressor of gene *i*.

This structure is represented numerically as a list of node structures whose fields represent the indices of its activators and repressors. This representation defines the set of ordinary differential equations used to simulate to the gene regulatory network dynamics.

The equations Eq. (6) were simulated for every gene through an Euler-Murayama scheme with a time step of *dt* = 0.001. In all simulations, we set *F* = 1 without loss of generality and *δF* = 1.0 to ensure a baseline production of proteins.

The transition timing was defined as the point at which the average activity of cluster 4 genes reached 95% of its maximum during the dynamics.

### Identification of parameters for the mathematical model

To simulate the gene expression dynamics, we sought to identify the optimal set of parameters in the e.o.m. that best captured the measured changes in gene activity, as calculated from the min-max normalization procedure.

Our mathematical model relies on linear regression beta coefficients as quantitative measures of interaction strength between genes. However, these beta-values do not directly correspond to gene activities. Firstly, the pluripotent state transition itself is a nonlinear process, meaning that the extent to which beta-values capture the strength of interactions between two genes depends on the specific cell clusters in which they are expressed. Additionally, beta-values may be influenced by other factors at the gene level, as gene expression dynamics result not only from direct interactions between two given genes but also from regulatory effects exerted by other genes. To address these challenges, we introduced additional parameters into our mathematical model to refine interaction strength at two levels: (1) between gene clusters (intercluster coefficients) and (2) between specific gene pairs.

For this, we assigned coefficients to gene interactions based on the specific combinations of clusters to which the genes belong (clusters 1 to 5, corresponding to naïve, early and later partially primed, early and late primed, respectively).

These coefficients scaled the gene interaction parameters by a numerical factor specific to each cluster pair. The choice of parameters was guided by the inferred GRN structure, with the pluripotent state transition progresses through sequential activation of clusters as follows: cluster 1 activates cluster 2, cluster 2 activates cluster 3, cluster 3 activates clusters 4&5. Additionally, clusters 4&5 self-reinforce, and all clusters exhibit mutual repression (as described in the Results section).

The intercluster interaction coefficients were manually optimised to reproduce the observed average cluster dynamics in RNAseq. As a result, activating interactions from cluster 3 to cluster 4 and 5, as well as the self-activation of cluster 4&5, were enhanced. Repressive interactions between cluster 1 and clusters 4&5 were adjusted to match the observed average cluster dynamics. Additionally, the incoming connections of genes that substantially deviated from their observed dynamics (*HNF4G*, *TGFB3*, *BMP2*, *PBX3*) were further refined.

This identified set of parameters was used to simulate the human pluripotency gene regulatory network with human and mouse expression data as inputs (see below), as well as for *in silico* knockouts.

### Machine learning-based optimisation of the network coefficients

To achieve the best reproduction of the dynamics observed in hPSCs, we optimised the network coefficients using the SciMLSensitivity software package in Julia, which is designed for fitting systems of differential equations parameters^95^.

First, we first applied min-max normalisation to the RNAseq expression values as follows:

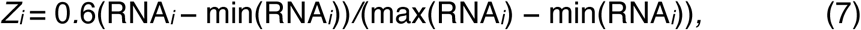

where RNA*_i_* are the expression level of gene *i*, and *Z_i_* is the experimentally obtained gene activity.

As such, min-max normalisation was performed within the range [0, 0.6], to allow for an increase in gene activity relative to wild-type dynamics, enabling simulations of gene perturbations (see below). The initial conditions for simulations were set to the obtained activities at the first measured time point (day 0). To accommodate the decrease in maximal gene activity, we also set *δF=* 0.85.

Next, we aimed to optimise the parameters of the model. Given the network complexity, manual tuning of parameters is impractical, while fully automated optimisation risks inconsistency due to the high degeneracy of the solution space given by the large number of parameters. To overcome this challenge, we combined manual and automatic approaches and introduced constraints aligned with the inferred network logic.

The parameters obtained from the manual approach described in “Identification of parameters for the mathematical model” section served as a starting point for subsequent automatic optimisation. The e.o.m parameters were then fitted using the SciMLSensitivity software package in Julia^95^, which is designed for parameter optimisation for a minimisation problem. We minimised the chi-squared difference between the observed experimental activities *Z_i_* and the simulated ones *z_i_* at the experimental time points:

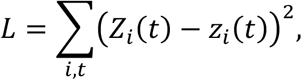

Minimization was obtained through the ParticleSwarm algorithm provided by the Optimization.jl library in Julia as follows. First, we optimized intercluster interactions with 140 particles for 100 iterations, adjusting our initial manually-chosen intercluster parameters. For consistency with the inferred network logic described above, we limited the optimum search to the interval [0.9,1.1] for all intercluster interactions, except for interactions to clusters 4&5 and repressive interactions from cluster 4&5 to clusters 1, 2, and 3, which were limited to the interval [0.1,30.0]. The resulting set of intercluster interactions parameters served as initial starting point for further optimization procedures.

Next, we optimized intergenic interactions for the 10 genes with the highest chi-squared difference between the simulated and observed dynamics, with constraints within an interval of width 80% of the previously obtained optimization result. This optimization has been done with the ParticleSwarm algorithm with 50 particles for 100 iterations. We then repeated this procedure again for the ensuing 10 genes with the highest chi-squared difference, with search of the corresponding parameters in the interval [0.0001,10] for TFs and [0.0,2.0] for signalling molecules.

### Quantification of interactions within the GRN

To quantitatively assess the strength of interactions within the GRN, we calculated an index for each sender-receiver pair, separately for positive and negative regulatory connections. A sender could represent either a gene or a gene cluster, while a receiver represented a gene cluster. The index was determined as the number of outgoing interactions from a given sender to a given receiver, normalised by the total number of outgoing interactions from the sender and the number of genes in the receiver.

### Simulation of single-gene knockout experiments

For *in silico* perturbations, single-gene knockouts were implemented by setting the activity of the knockout gene at 0 throughout the entire simulation. We simulated the gene network using e.o.m. as described above, for a total time of *T* = 20. The transition time was defined as the point at which the average activity of cluster 4 genes reached 95% of its maximum during the dynamics. This defines the delay or acceleration of the knock-out perturbation relative to the wild-type control.

### Simulations of human pluripotency GRN using mouse data

Simulations of the human pluripotency GRN using mouse data as input were performed using three publicly available bulk RNAseq datasets from mouse pluripotent state transition *in vitro*^47–49^, and the e.o.m. described above. Where available, expression values from multiple replicates were averaged.

Gene activities from mouse RNAseq expression values were obtained by three different normalisation methods. In the “initial-time” normalisation, RNAseq expression values were min-max normalised over the initial time point (day 0). In the “time-course” normalisation, min-max normalisation was applied to each gene across the time-course. In the “total” normalisation, min-max normalisation was performed across all genes and time-points.

Simulations of mouse dynamics were run using the manually-chosen parameters (see above) over the human dynamics, with mouse data as initial conditions for different normalisation approaches. As all three normalisation methods produced consistent results (Table S7), only the total normalisation approach is presented in the Results section.

### Naïve hPSC transfection

Naïve hPSCs were transfected either chemically or using the NEON Transfection System (ThermoFisher Scientific), according to the manufacturers’ protocols.

For chemical transfection, 4.4μg of DNA were thoroughly mixed with 204μl of OptiMEM medium and 15.6μl of FuGENE reagent (Promega), followed by incubation for 20 minutes. 10^6^ naïve hPSCs were dissociated into single-cells using TrypLE, pelleted using centrifugation (3 minutes at 200g), and resuspended in the DNA solution. After 2 minutes of incubation, the cells were plated into one well of a 6-well plate with irradiated MEFs in PDLGX medium (see above) supplemented with 10μM ROCK inhibitor and 1μl/ml Geltrex, as during routine passaging. The medium was refreshed the next day.

For Neon electroporation, 1.2x10^6^ naïve hPSCs were dissociated to single-cells using TrypLE, pelleted using centrifugation (3 minutes at 200g) and resuspended in 120μl of Resuspension Buffer R from the NEON 100μl kit (Cat. MPK10096, ThermoFisher Scientific). The cells were then mixed with 9.6μg of DNA prepared at a concentration of 1-5μg/μl. Since the mix was prepared with a 20% excess, transfection was performed using 10^6^ cells and 8μg of DNA. The cells were aspirated into a 100μl NEON tip, which was then placed into a NEON tube containing 3ml of Electrolytic Buffer E2. Transfection was carried out using the following programme: 1,150V, 30ms, 2 pulses. The cells were plated into one well of a 6-well plate with irradiated MEFs in PDLGX medium supplemented with 10μM ROCK inhibitor and 1μl/ml Geltrex, as during routine passaging. Antibiotics, such as Penicillin-Streptomycin, should not be applied for 24h after transfection. The medium was refreshed the next day.

### Gene knockout in naïve hPSCs

For gene knockout, two gRNAs were designed to introduce cuts and cause a frameshift within an exon shared by as many transcripts of a gene of interest as possible. gRNAs were designed using the Wellcome Sanger Institute Genome Editing online tool (https://wge.stemcell.sanger.ac.uk/).

gRNAs were cloned into pSpCas9(BB)-2A-GFP (PX458) plasmids (a gift from Feng Zhang; Addgene plasmid # 48138; http://n2t.net/addgene:48138; RRID:Addgene_48138)^73^. gRNA sequences are listed in the Key Resource Table.

10^6^ naïve hPSCs were co-transfected with 4μg of each gRNA-encoding plasmid (totalling 8μg of transfected DNA) using NEON Transfection System. GFP-positive cells were flow sorted 48 hours post-transfection and plated at a concentration of about 100-200 cells/cm^2^.

For *ESRRB* and *TBX3* gene knockouts, clonally derived populations were established as described below. Since the gene knockout efficiency for these genes was approximately 90%, a different strategy was applied for *NFKB1* gene knockout. For *NFKB1*, the resulting clones were pooled for further maintenance and experimentation; gene downregulation was confirmed using qRT-PCR.

Single-cell derived clones were manually picked 12-16 days post-transfection, dissociated with TrypLE and plated into 96-well plates with irradiated MEFs.

After clone expansion, approximately 2x10^4^ cells were treated with 85-100μl of a non-ionic detergent-containing lysis buffer (NID buffer) containing 10mM Tris HCl pH8.0, 50mM KCl, 2mM Mg_2_Cl, 0.1mg/ml gelatin, 0.45% Tween-20, 0.45% NP40 and 120μg/ml Proteinase K, for 2 hours at 55 °C, followed by Proteinase K inactivation for 10 minutes at 95 °C. PCR genotyping was performed using 2μl of the cell lysate.

Deletions were sequence-verified in selected clones. For this, the targeted genomic regions were PCR amplified using Taq polymerase (M0267, NEB), extracted from a gel (Monarch DNA Gel Extraction kit, NEB) and analysed using standard Sanger sequencing (Genewiz, Azenta).

To validate the diploid state, cells were fixed using 70% ethanol, followed by staining with 50μg/ml propidium iodide in the presence of 0.5mg/ml RNase A. DNA content was measured using flow cytometry and compared to a diploid control.

### Gene knockdown in naïve hPSCs

For gene knockdown, three gRNAs were designed to target as many transcripts of a gene of interest as possible, ensuring they recognise a target sequence within 100nt downstream of the TSS. gRNAs were designed either CRISPick (https://portals.broadinstitute.org/gppx/crispick/public) or CHOPCHOP tools (https://chopchop.cbu.uib.no/). gRNA sequences are listed in the Key Resource Table.

Gene knockdown was carried out as previously described^74^ using the constructs reported in this study (see below).

gRNAs were cloned into pPB-Ins-U6p-sgRNAentry-EF1Ap-TetOn3G-IRES-Neo plasmid (a gift from Azim Surani; Addgene plasmid #183411; http://n2t.net/addgene:183411; RRID:Addgene_183411).

Gene knockdown was achieved using an inducible catalytically-dead Cas9 (dCas9) fused with the KRAB domain delivered through the CRISPRi plasmid (pPB-Ins-TRE3Gp-KRAB-dCas9-ecDHFR-IRES-GFP-EF1Ap-Puro, a gift from Azim Surani; Addgene plasmid #183410; http://n2t.net/addgene:183410; RRID:Addgene_183410).

The above constructs were stably integrated into the genome of naïve hPSCs using the hyperactive PiggyBac transposase (hyPBase) delivered through pCMV-hyPBase plasmid (a gift from Allan Bradley^122^) 10^6^ naïve hPSCs were co-transfected with 1.466μg of plasmids containing cloned gRNAs (in equal proportions); 1.466μg of CRISPRi plasmid and 1.466μg of pCMV-hyPBase (totalling 4.4μg of transfected DNA) using FuGENE, as described above. Selection was started with 50μg/ml geneticin (G418) and 0.2μg/ml puromycin 48 hours post-transfection to obtain stably transfected cells. Antibiotic concentrations were gradually increased over 14 days to 200μg/ml G418 and 0.5μg/ml puromycin. The stably transfected clones were pooled by passaging using TrypLE. The resulting cell population was routinely maintained in the presence of 200μg/ml G418 and 0.5μg/ml.

To test the effect of genetic knockdown during the pluripotent state transition, 1μg/ml doxycycline and 10μM Trimethoprim were added 48h prior to the transition (otherwise performed as described above). Cell pellets from the induced and not induced cells were collected at different time points during the transition.

### qRT-PCR

Total RNA was extracted using RNeasy Mini Kit (74104, Qiagen) and 500ng was used for reverse transcription using RevertAid First Strand cDNA Synthesis kit (ThermoFisher Scientific). Quantitative PCR was performed with Brilliant III Ultra-Fast SYBR Green qRT-PCR Master Mix (Agilent). Primer sequences are listed in the Key Resource Table.

## QUANTIFICATION AND STATISTICAL ANALYSIS

10X Multiomics was done in 1 biological replicate per time point, aiming to sequence 10,000 cells per sample. Five pseudobulk replicates per cell cluster were created by bootstrapping 30% of cells for differential expression and accessibility analyses.

Differential expression and accessibility analysis were conducted using the following thresholds: FDR <0.01, abs(log2 Fold Change) ≥1.

For gene set and pathway overrepresentation analyses, genes of interest were compared against a background of all annotated genes (n = 36,601), using a significance threshold of FDR <0.05.

Metacells were inferred using SEACells, converging after 21 iterations (epsilon = 1e-4) and yielding 316 metacells (1.3% of total cells), of which 250 (median 85.5 cells per metacell) were retained after QC. Metacells were used for the GRN inference.

For the *in silico* ChIP, peaks within 50kb of the gene body were considered potential regulatory elements for the respective gene. TF binding was defined by an *in silico* ChIP score >0.06.

Activating and repressing interactions within the GRN were identified using linear regression, with metacell expression values of the sender as an independent variable and those of the receiver as a dependent variable. Significant interactions were defined by abs(beta) >0.24 and p-value < 0.1.

For signaling pathways, regulatory connections were considered significant if they met the following criteria: (1) weight of connection >0.1 from NicheNetR; (2) abs(beta) >0.24 and p-value < 0.1 from linear regression for ‘ligand-target’; (3) receptor changes in the same direction as the respective ligand or does not significantly change.

For mathematical modelling, the gene expression values were scaled between [0,1], with mRNA assumed to be proportional to protein levels and a degradation rate in human *b*=1. Simulations were run with a time step of *dt* = 0.001 for a total of *T*=20; assuming *F* = 1 and *δF* = 1.0. The transition time was defined as the point at which the average activity of cluster 4 genes reached 95% of its maximum during the dynamics. The similarity of simulated gene expression dynamics and RNAseq was assessed using a chi-squared test, with a random network used for comparison.

For model optimisation, gene expression values were scaled between [0,6] and *δF* was set to 0.85. ML-based automated optimisation was first applied to intercluster interaction parameters, followed by optimisation of intergenic interaction parameters for genes with the greatest differences between simulated and observed dynamics.

For simulations of human GRN using mouse TF levels as the initial input, gene activities from mouse RNAseq expression values were obtained by three different normalisation methods: (1) min-max normalisation over the initial time point (day 0); (2) min-max normalisation across the time-course for each gene; (3) min-max normalisation across all genes and time-points. Protein turnover in mice was considered to be 2.3 times faster than in human, based on existing data.

qRT-PCR error bars were derived from two technical replicates in Figures 4H, S4D, S6C, S6D; or for two biological replicates (that is two independent transition experiments) in Figures 4J.

qRT-PCR was performed for 3 biological replicates in Figure S4 (that is three independent transition experiments).

